# Phosphorylation enables allosteric control of a viral condensate

**DOI:** 10.1101/2025.05.24.655949

**Authors:** Julia Acker, Xinyu Wang, Daniel Desirò, Tanushree Agarwal, Alice Colyer, Cyril Haller, Rob Scrutton, Lee Sherry, Kadi L Saar, Rosie Murray, Ksenia Fominykh, Sai Hou Chong, Jeremy D. Schmit, Antonio N. Calabrese, Tuomas P. J. Knowles, Alexander Borodavka

**Author notes:** These authors contributed equally to this work (Julia Acker, Xinyu Wang). School of Infection & Immunity, University of Glasgow, Glasgow, UK.

## Abstract

In many viruses, intrinsically disordered proteins (IDPs) drive the formation of replicative organelles essential for viral production. In species A rotaviruses, the disordered protein NSP5 forms condensates in cells via liquid-liquid phase separation (LLPS). Yet the sequence diversity of NSP5 raises the question of whether condensate formation is conserved across all strains and if distinct variants employ alternative mechanisms for nucleating phase separation. Using a machine learning approach, we demonstrate that NSP5 variants differ significantly in their propensity to phase-separate. We engineered a variant incorporating amino acid signatures from strains with low LLPS tendency, which failed to phase separate in vitro yet supported the formation of replicative condensates in recombinant viruses in cells. Low-tendency LLPS strains require phosphorylation of NSP5 to nucleate phase separation, whereas high-tendency strains do not, suggesting distinct nucleation mechanisms. Furthermore, hydrogen-deuterium exchange mass spectrometry revealed a phosphorylation-driven allosteric switch between binding sites on the high-propensity variant. These findings establish that phosphorylation plays a context-dependent role in the formation of replicative organelles across diverse rotaviruses.

## Introduction

Liquid-liquid phase separation (LLPS) underpins the formation of cellular compartments lacking a membrane (Hyman et al., 2014) including P-bodies, stress granules, and nucleoli (Banani et al., 2017; Brangwynne et al., 2011, 2009). LLPS is driven by weak interactions between multivalent, polymer-like molecules (Banani et al., 2017; Brangwynne et al., 2015). This mechanism is commonly explained using the “stickers and spacers” model (Choi et al., 2020), where “stickers” refer either to side chain interactions within intrinsically disordered regions (IDRs) (Martin et al., 2020; Wang et al., 2018) or to interactions involving folded domains that recognise specific binding motifs (Banani et al., 2016). Regions separating the stickers, known as spacers, modulate interactions mediated by the stickers (Ginell and Holehouse, 2023). The hub-and-driver model describes a system in which a “hub” molecule characterised by a low intrinsic propensity for LLPS binds to “driver” molecules whose IDRs confer a higher LLPS propensity (Galagedera et al., 2023).

While LLPS is predominantly governed by inherent physico-chemical characteristics of macromolecules, understanding the emergent properties of condensates presents a formidable task. Due to their minimalistic proteomes, viruses provide excellent models for dissecting the roles of a relatively small number of components that drive LLPS, and indeed, many viruses employ LLPS to form viral factories that support their replication (Borodavka and Acker, 2023; Heinrich et al., 2018; Monette et al., 2020; Nikolic et al., 2016; Zhou et al., 2022).

Group A rotaviruses (RVA) represent a large and diverse class of important double-stranded RNA pathogens that infect humans and other animals worldwide. Several RVA strains that infect mammals, including simian SA11 and RRV, bovine RF, and porcine OSU have all been shown to form cytoplasmic replication factories, termed viroplasms (Campagna et al., 2007; Carrenno-Torres et al., 2010; Geiger et al., 2021). Viroplasms represent protein-RNA condensates formed via LLPS of the non-structural protein NSP5 and the RNA chaperone NSP2 (Geiger et al., 2021). While both proteins are required for nucleating phase separation *in vitro*, NSP5 is believed to be the LLPS scaffold/driver, and NSP2 is a major client/hub (Geiger et al., 2021; Nichols et al., 2023). The onset of viroplasm formation depends on the expression levels of both proteins, and on the multiplicity of infection and its duration (Geiger et al., 2021). The 35 kDa RNA chaperone NSP2 forms an octamer with positively charged grooves involved in NSP5/RNA binding (Bravo et al., 2021; Jayaram et al., 2002; Jiang et al., 2006). In contrast, NSP5 is an intrinsically disordered protein (IDP) harbouring an 18 amino acid C-terminal homo-oligomerisation region (CTR) (Geiger et al., 2021). Removal of the CTR renders the remaining IDR unable to drive LLPS when mixed with NSP2 *in vitro* (Geiger et al., 2021). Amino acids 1 - 33 and the CTR of NSP5 are involved in binding to NSP2 (Arnoldi et al., 2007; Eichwald et al., 2004b; Fabbretti et al., 1999). During infection, the 21 kDa NSP5 undergoes a phosphorylation cascade leading to the formation of phospho-isoforms ranging from 26 kDa to 35 kDa (Afrikanova et al., 1998; Eichwald et al., 2004a). In the cell-culture adapted strain SA11, nine phosphorylation sites, including Ser2/4, Ser30, Ser37, Ser42, Ser56, Ser67, Ser101, Ser127, and Ser164, have been reported (Sotelo et al., 2010). These residues undergo phosphorylation following the initial phosphorylation of the conserved Ser67 upon NSP2/NSP5 binding (Papa et al., 2019). It remains unclear how sequential NSP5 phosphorylation influences phase separation and the functional state of viroplasms across diverse RVA variants.

Phosphorylation can modulate the phase behavior of IDPs in diverse ways. It may enhance liquid-like properties in condensates formed by the Adenovirus L1-52/55 kDa protein (Grams et al., 2024), or protect against aberrant liquid-to-solid transitions (Ranganathan et al., 2023). Conversely, phosphorylation may reduce condensate formation (Bressler et al., 2023), highlighting the system-dependent nature of phosphoregulation. Mechanistically, phosphorylation has been proposed to alter the conformational ensemble of IDPs, whereby it can influence subsequent interactions or modifications within the same disordered polypeptide chain through a process known as dynamic allostery (Berlow et al., 2018; Ferreon et al., 2013; Hilser and Thompson, 2007; Kern and Zuiderweg, 2003) However, identification of such allosteric events within IDPs that drive changes in the architecture and composition of condensates remains challenging.

It remains unclear whether all RVA strains utilise LLPS, and if they do, whether they rely on a common, conserved mechanism. Given the large number of RVA strains encoding diverse NSP5 sequences, assessing their individual propensities for LLPS is challenging. This difficulty is compounded by the limited ability of many RVA isolates to grow in cell culture, with strains SA11 and RF being the most characterised and readily grown in cell culture (Arnold et al., 2012; Pena-Gil et al., 2023; Ward et al., 1984). To overcome these limitations, we employed a machine learning algorithm (Saar et al., 2021) to evaluate the phase separation potential of various NSP5 variants and engineered an SA11-like strain expressing an NSP5 variant with a low LLPS score, designated SC_low_.

Our comparative analysis of a low-tendency LLPS variant alongside high-tendency variants reveals that phosphorylation plays a dual role: it nucleates phase separation in low-propensity NSP5 variants and modulates binding interactions within condensates in high-propensity variants. This dual function suggests that the capacity for NSP5 to undergo phase separation is a conserved feature essential to the replication cycle of RVAs, with low-scoring variants acquiring this capability through phosphorylation, underscoring phosphorylation as a pivotal regulator of viroplasm assembly.

## Results

### Engineering an NSP5 variant with diminished LLPS potential

To evaluate the condensate-forming potential of NSP5 across a range of RVA strains and isolates, we performed an *in silico* analysis of their LLPS propensities using DeePhase (Saar et al., 2021). Of the 451 unique RVA NSP5 sequences retrieved from the NCBI virus database, most exhibited DeePhase scores above 0.5 (**Fig. 1A**), indicating a high propensity for LLPS (Saar et al., 2021). Notably, the NSP5 protein from the cell culture-adapted SA11 strain exhibited the highest DeePhase score among all sequences analysed. Seven NSP5 variants designated S1_low_ to S7_low_ displayed DeePhase scores below 0.4 (**Table 1**), suggesting a limited or absent ability to undergo phase separation.

**Figure 1.**
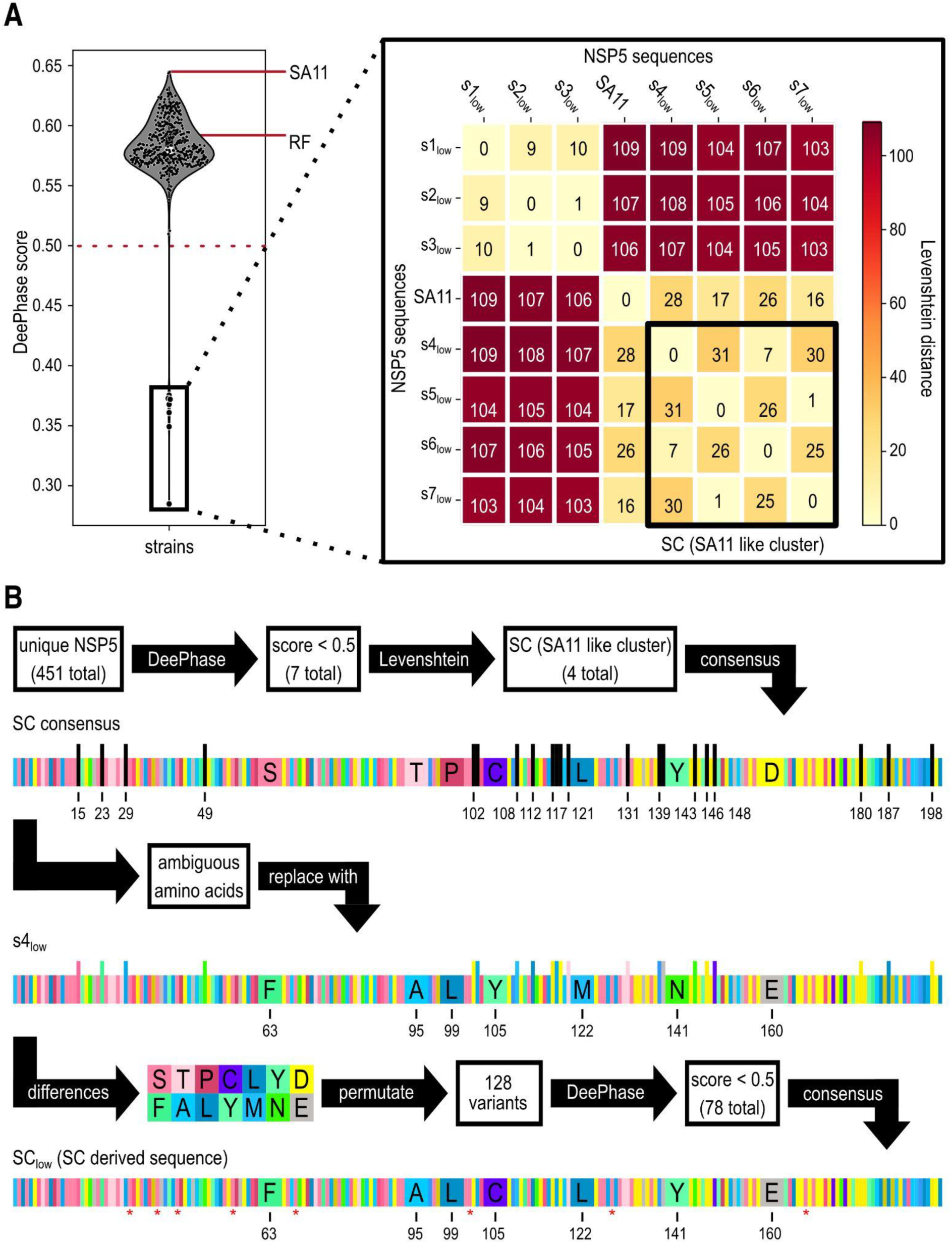
Computational identification and rational design of an NSP5 variant with reduced LLPS propensity. **A.** LLPS propensity analysis of 451 unique full-length RVA NSP5 sequences using DeePhase. The violin plot shows that most NSP5 variants exhibit DeePhase scores above 0.5. The cell-culture adapted RVA strains RF and SA11 highlighted. A zoomed-in inset displays the seven low-scoring variants designated S1_low_-S7_low_. Sequence similarity was assessed by computing pairwise Levenshtein distances, ranging from 0 (identical sequence) to 110 (maximally divergent). Among these, NSP5 sequences from S4_low_ - S7_low_ show the greatest similarity with strain SA11 and are collectively referred to as the SA11-like cluster (SC). **B.** *In silico* design of the SC_low_ NSP5 variant using DeePhase predictions and NSP5 conservation analyses. A consensus sequence (SC consensus) was generated from the SA11-like cluster, incorporating 21 degenerate positions, which were substituted with corresponding residues from the S4_low_ sequence. The seven remaining differences between the SC consensus and S4_low_ gave rise to 2^7 (128) sequence permutations, each evaluated by DeePhase. Of these, 78 variants scored below 0.5. A second consensus was derived from these low-scoring variants, resolving one remaining ambiguous position by adopting the S4_low_ amino acid. Asterisks mark serine phosphorylation sites reported for NSP5 (Sotelo et al., 2010), all of which are retained in the final SC_low_ sequence.

**Table 1.**
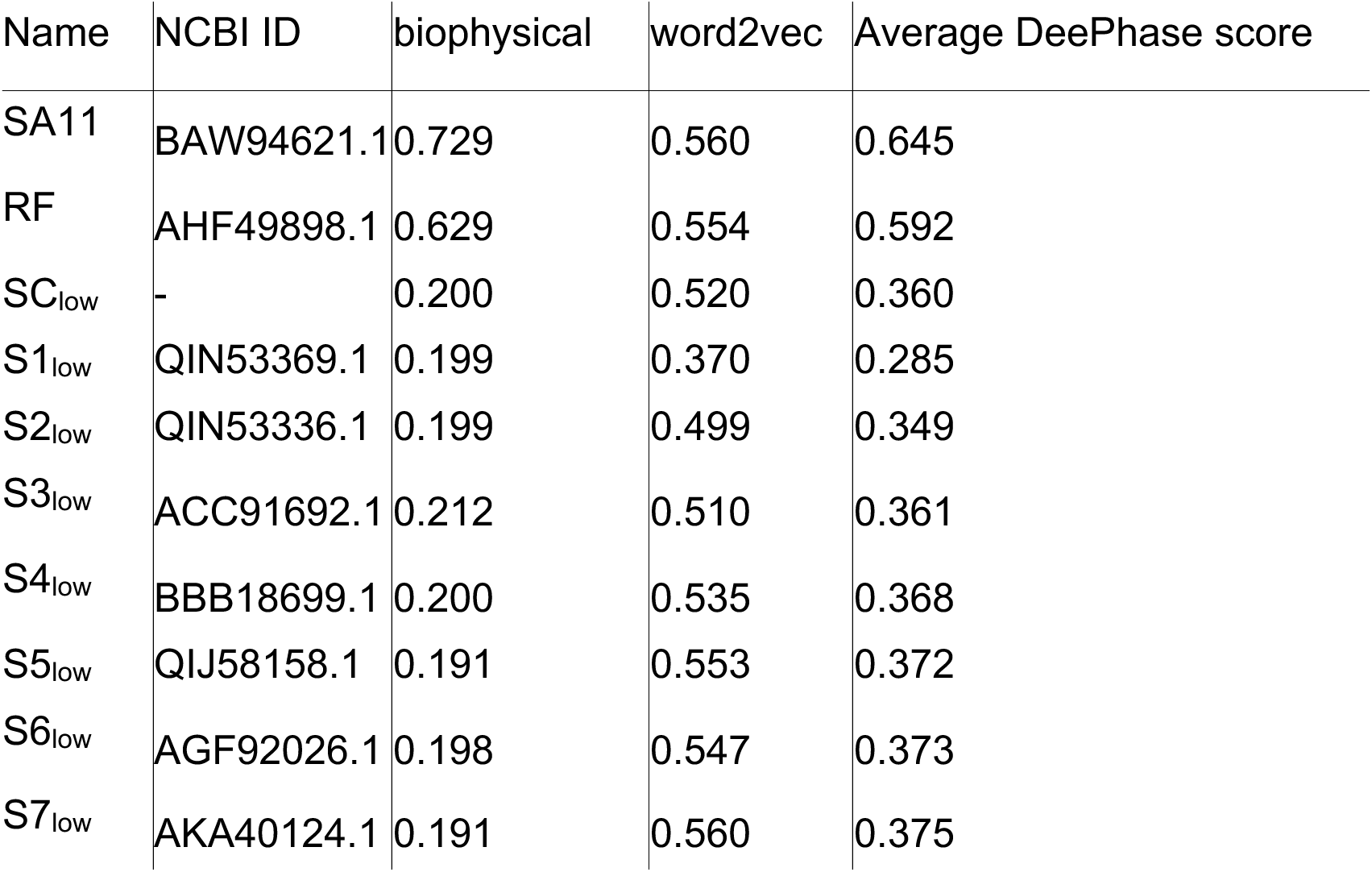
The DeePhase (Saar et al., 2021) scores for all major sequences used in this work are listed in the propensity column. All scores were calculated from the average of the biophysical and word2vec columns. The name of each sequence referred to here is listed in the name column. The NCBI ID column, on the other hand, lists the sequence-specific ID under which it can be found in the NCBI GenBank. In addition, the length column indicates the number of amino acids in each protein sequence.

To estimate sequence divergence between the well-characterised reference strain SA11 and the identified low-propensity NSP5 variants, we calculated pairwise Levenshtein distances between each S1_low_ to S7_low_ and SA11 sequences. The Levenshtein distance quantifies the minimum number of single-residue changes needed to transform one sequence into another, providing a straightforward numerical measure of their overall similarity (Levenshtein, 1966). The analysis identified four SA11-like variants, designated S4_low_ through S7_low_ that form an SA11-like cluster (SC) based on their shortest pairwise Levenshtein distances to each other and to SA11 (**Fig. 1A**).

We then engineered an SA11-like NSP5 variant with reduced LLPS propensity by aligning the four naturally occurring SC variants (S4_low_ - S7_low_) (Katoh and Standley, 2013a), and deriving a consensus sequence that maintained NSP5 conserved regions while introducing only amino acid substitutions predicted to reduce its LLPS score. Given the 21 degenerate positions in the SC variants (**Fig. 2B**), we replaced each of these residues with the corresponding amino acid from the S4_low_ variant, which exhibited the lowest DeePhase score (**Table 1**). Since the resulting sequence diverged from S4_low_ at seven amino acid positions, we generated all 2^7, or 128 possible combinations of these substitutions and evaluated each variant using DeePhase.

**Figure 2.**
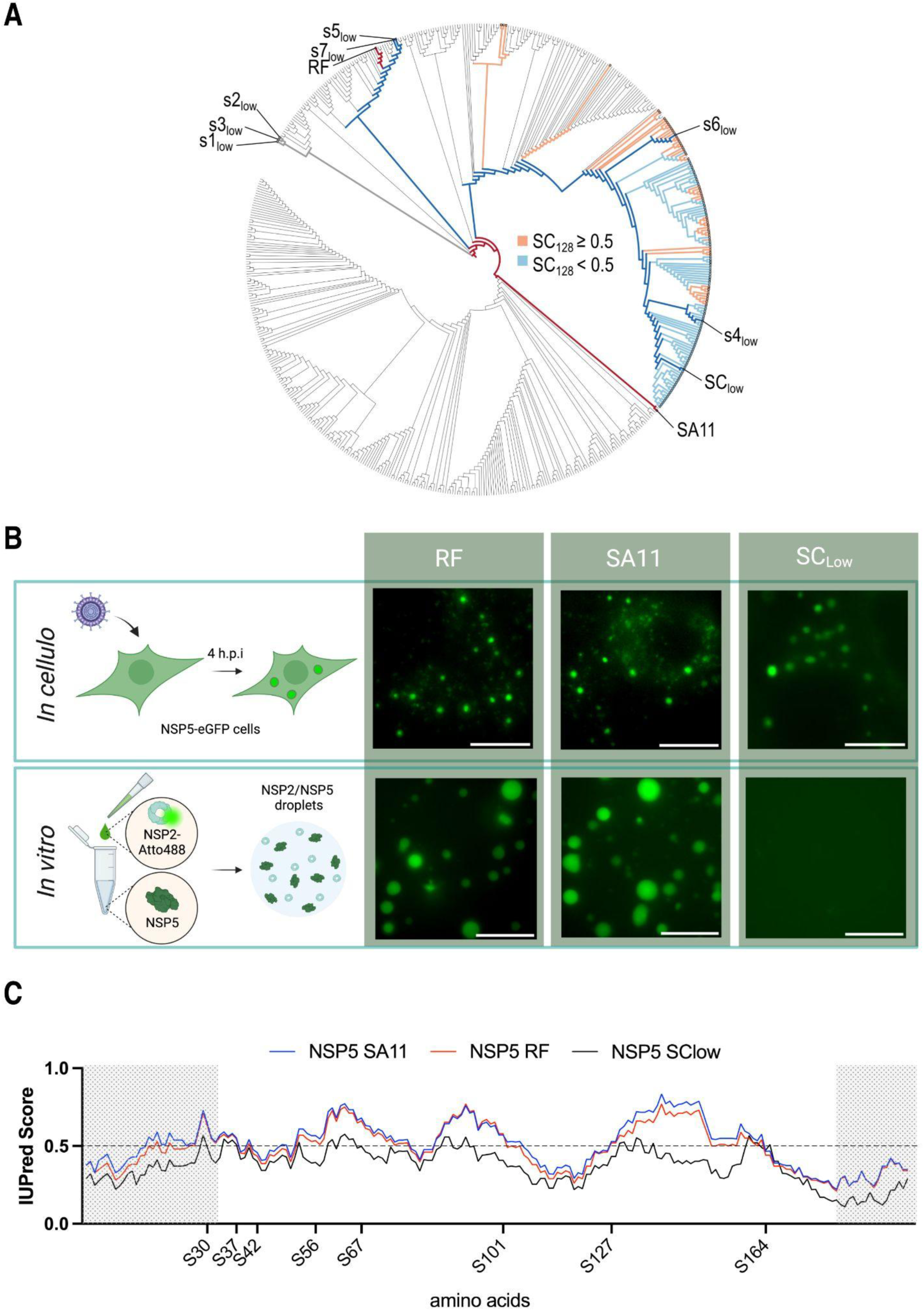
Phylogenetic and biophysical features underlying NSP5 phase separation propensity. **A.** Phylogenetic analysis of 451 unique NSP5 sequences alongside 128 computationally designed permutation variants derived from the SA11-like cluster. Naturally occurring low-scoring variants (S1_low_-S7_low_; DeePhase < 0.5) are indicated at their respective tips. Cell-culture-adapted reference strains SA11 and RF are highlighted in dark red, and the SA11-like cluster (SC) consensus with its optimised SC_low_ variant are marked in dark blue. Among the 128 permutation variants (SC128), tips are coloured orange for sequences with DeePhase scores > 0.5, while light blue denotes scores < 0.5. **B.** NSP2–NSP5 condensate formation in model RVA strains RF and SA11, observed both *in cellulo* and *in vitro*. *Upper panel*: MA104–NSP5–eGFP cells infected with either RF or SA11 RVAs develop cytoplasmic condensates. *Lower panel*: recombinantly purified and Atto488-labelled NSP5 and NSP2 from RF and SA11 strains form condensates *in vitro*. Scale bars, 10 µm. **C.** Intrinsic disorder predictions for NSP5 from RVA strains SA11 (blue) and RF (red), as determined by AUPred2A (Mészáros et al., 2018). Scores above 0.5 (dashed line) indicate disordered regions. The N-terminal (residues 1–33) and C-terminal (residues 181–198) segments (shaded) are required for NSP2 binding and NSP5 homo-oligomerisation, respectively (Arnoldi et al., 2007; Eichwald et al., 2004b; Fabbretti et al., 1999). Phosphorylated serine residues reported by Sotelo *et al*. (Sotelo et al., 2010) are marked.

Seventy-eight of the 128 variants scored below 0.5 (**Fig. 1B**), demonstrating that substitutions at these seven positions significantly influence the DeePhase score of the SA11-like SC variants (**Fig. 1B and 2A**). Notably, all amino acid substitutions mapped exclusively to the predicted intrinsically disordered region (IDR) of NSP5, suggesting that alterations within the IDR alone are sufficient to differentiate low-from high-LLPS-propensity NSP5 variants. We then generated a consensus sequence using the 78 low-LLPS score NSP5 variants with a single residue remaining ambiguous. This ambiguity was resolved by selecting the corresponding amino acid from S4_low_ variant to generate the final SC_low_ sequence that achieved the DeePhase score of 0.36.

### A low-LLPS NSP5 variant supports viroplasm formation and viral replication

We next evaluated whether the SC_low_ NSP5 variant could support replication of an SA11-like RVA in cell culture. Given the segmented nature of the RVA genome, we considered the possibility that sequence mismatches between the SC_low_ NSP5 gene (gene segment 11) and the SA11 backbone might impair virus rescue or propagation. To minimise potential inter-segmental incompatibility, we synonymously substituted all nucleotides in the SC_low_ NSP5 coding sequence that differed from the SA11 reference, selecting synonymous codons with the smallest evolutionary distance to the original sequence, as detailed in the Methods (**Expanded view Fig 3A**).

**Figure 3.**
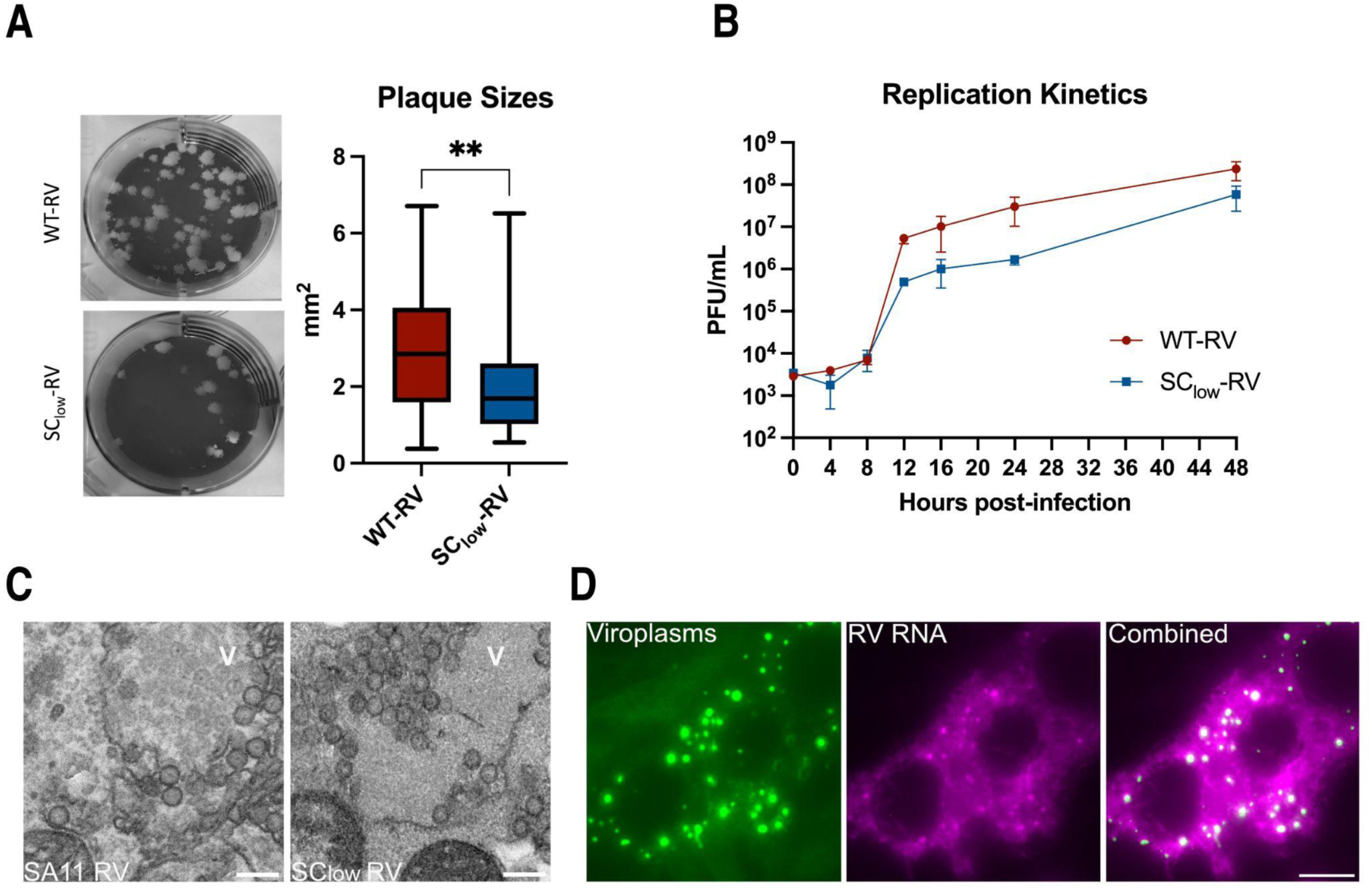
SC_low_-RV supports replication, particle production, and viroplasm formation in cell culture. **A.** Representative plaque morphologies produced by SC_low_-RV and and wild-type SA11-RV in MA104 cells. Box plots show plaque areas measured for 50 individual plaques. The horizontal bar indicates the mean ± standard error of the mean (SEM). **p = 0.0033, unpaired t-test. **B.** Replication kinetics of SC_low_-RV and wild-type SA11-RV in MA104 cells infected at a multiplicity of infection (MOI) of 1. Viral titres were determined at 4, 8, 12, 16, 24, and 48 hours post infection (h.p.i.) using TCID₅₀ assays. Each data point represents the mean ± standard deviation (SD) from three independent experiments. **C.** Transmission electron micrographs of MA104 cells infected with SA11-RV (left) and SC_low_-RV (right), fixed at 8 h.p.i. Multiple viral particles are observed within viroplasms (V). Scale bar, 200 nm. **D.** SC_low_-RV forms viroplasms that accumulate viral transcripts. Atto647N-labelled smFISH probes specific for RV transcripts were used to visualise viral RNAs localising to EGFP-tagged viroplasms, as described in Methods.

**Expanded view Figure 3.**
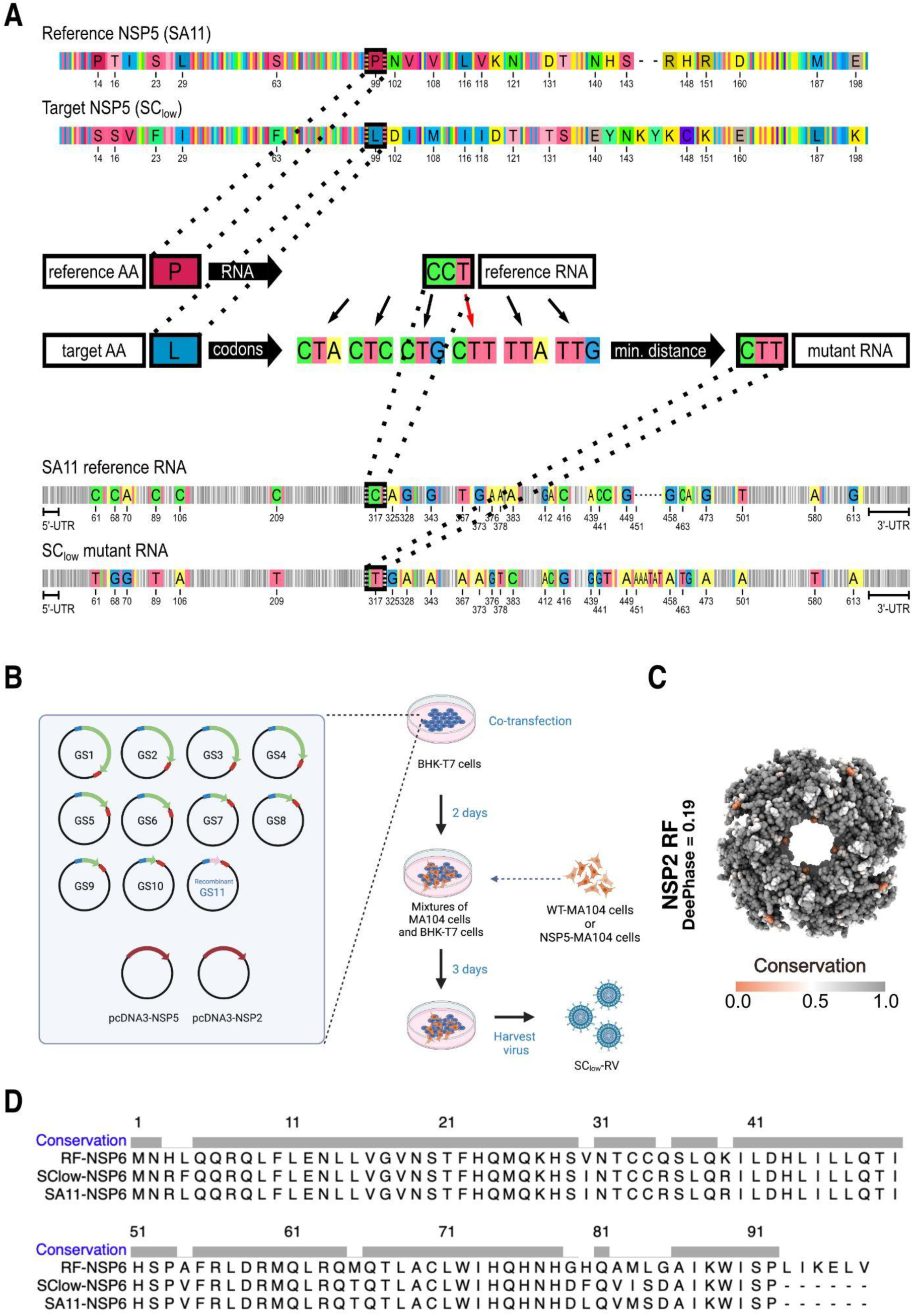
Design, rescue, and genetic stability of the SC_low_ rotavirus. **A.** RNA sequence design for gene segment 11 SC_low_ ORF using the reference RNA sequence of NSP5 SA11. The coding sequence of NSP5 from SA11 was used as a reference, and synonymous codons were selected to minimise nucleotide divergence while preserving amino acid identity wherever possible. In cases of amino acid substitution (e.g., Proline to Leucine at position 99), the target codon required the smallest possible number of nucleotide changes. A total of 35 nucleotide changes were introduced in the SC_low_ sequence, while untranslated regions (UTRs) were left unchanged. Positions of nucleotide substitutions are indicated. **B.** Schematic overview of the reverse genetics strategy used to generate SC_low_-RV. Ten pT7 plasmids encoding the SA11 gene segments (GS1–GS10) and one recombinant plasmid encoding SC_low_, were co-transfected into BHK-T7 cells along with expression plasmids for NSP2 and NSP5 (pcDNA3-NSP2 and pcDNA3-NSP5). At 48 h post-transfection, either WT-MA104 or NSP5-expressing MA104 (NSP5-MA104) cells were overlaid. Recombinant virus was harvested after three freeze-thaw cycles upon observation of full cytopathic effect (CPE), as described in Methods (Papa et al., 2019). **C.** Conservation analysis of NSP2 across rotavirus A strains. Full-length NSP2 sequences were aligned, and residue conservation was mapped onto the NSP2 octamer crystal structure (PDB: 1L9V; SA11 strain). Key NSP5-binding regions are conserved across strains (Jiang et al., 2006). **D.** Multiple sequence alignment of the NSP6 open reading frame encoded by gene segment 11. The NSP6 sequences of SA11, RF, and SC_low_ sequence were compared, confirming preservation of NSP6 in the engineered SC_low_-RV genome.

We used a modified plasmid-based reverse genetics system (**Expanded view Fig 3B**), incorporating the SC_low_-encoding gene segment 11, to rescue SC_low_-RV in an MA104 cell line stably expressing wild-type NSP5 (NSP5-MA104), a platform previously employed to recover NSP5-deficient viruses (Papa et al., 2019). Upon rescue, SC_low_-RV could also replicate in wild-type MA104 cells, confirming that SC_low_ NSP5 was functional. Sequencing of the SC_low_ NSP5 gene from recombinant rotavirus indicated that its sequence remained stable for at least ten passages in MA104 cells at a low multiplicity of infection **(Supplementary File).**

Regions of NSP2 previously implicated in NSP5 binding (residues 64–68, 179–183, 232–251 and 291–302 (Jiang et al., 2006)) were conserved in SC_low_-RV and remained unchanged after ten sequential passages, suggesting that NSP2 retains its ability to interact with the low-scoring NSP5 variant (**Expanded view Fig 3C**). As SA11 gene segment 11 also encodes NSP6, which enhances viral growth (Komoto et al., 2017), we analysed the modified segment and confirmed that NSP6 was retained in SC_low_-RV (**Expanded view Fig 3D**).

To further characterise SC_low_-RV, we compared plaque morphologies of SC_low_-RV and SA11-RV. Despite comparable end-titres, plaques formed by SC_low_-RV were significantly smaller than those of SA11-RV (2.0 ± 1.4 mm^2^ vs 3.0 ± 1.7 mm^2^, respectively) (**Fig 3A)**. We next assessed SC_low_-RV by comparing its replication kinetics with the WT virus in MA104 cells infected at a multiplicity of infection (MOI) of 1. SC_low_-RV replicated more slowly than SA11-RV (**Fig 3B**), although this difference diminished at later stages of infection (48 h.p.i), consistent with the similar end-titres observed. We further confirmed viral particle production in viroplasms formed by SC_low_-RV using electron microscopy (EM) imaging at 8 h.p.i, which revealed the presence of multiple viral particles adjacent to viroplasms (**Fig 3C**). Finally, single-molecule RNA Fluorescence In Situ Hybridisation (smFISH) (Strauss et al., 2023), confirmed that viroplasms formed by SC_low_-RV harboured viral transcripts (**Fig 3D**). Taken together, we show that SC_low_-RV supports viroplasm formation and robust replication in cell culture.

### SC_low_-RV displays delayed viroplasm maturation and reduced NSP5 hyperphosphorylation

Having established that SC_low_-RV replicates and forms viroplasms in infected cells, we next examined the kinetics of viroplasm formation in cells infected with SC_low_-RV compared to SA11-RV. To visualise viroplasms, we infected NSP5-eGFP MA104 cells with either SC_low_-RV or SA11-RV at an MOI of 20. Infected cells were imaged at 4, 6, and 8 hours post infection (h.p.i.). Both SC_low_-RV and SA11-RV produced spherical NSP5–eGFP-tagged condensates as early as 4 h.p.i. (**Fig 4A)**. However, a marked reduction in both the abundance and size of viroplasms was observed in SC_low_-RV infected cells compared to SA11-infected cells at early time points (up to 8 h.p.i.) (**Fig 4B & C**).

**Figure 4.**
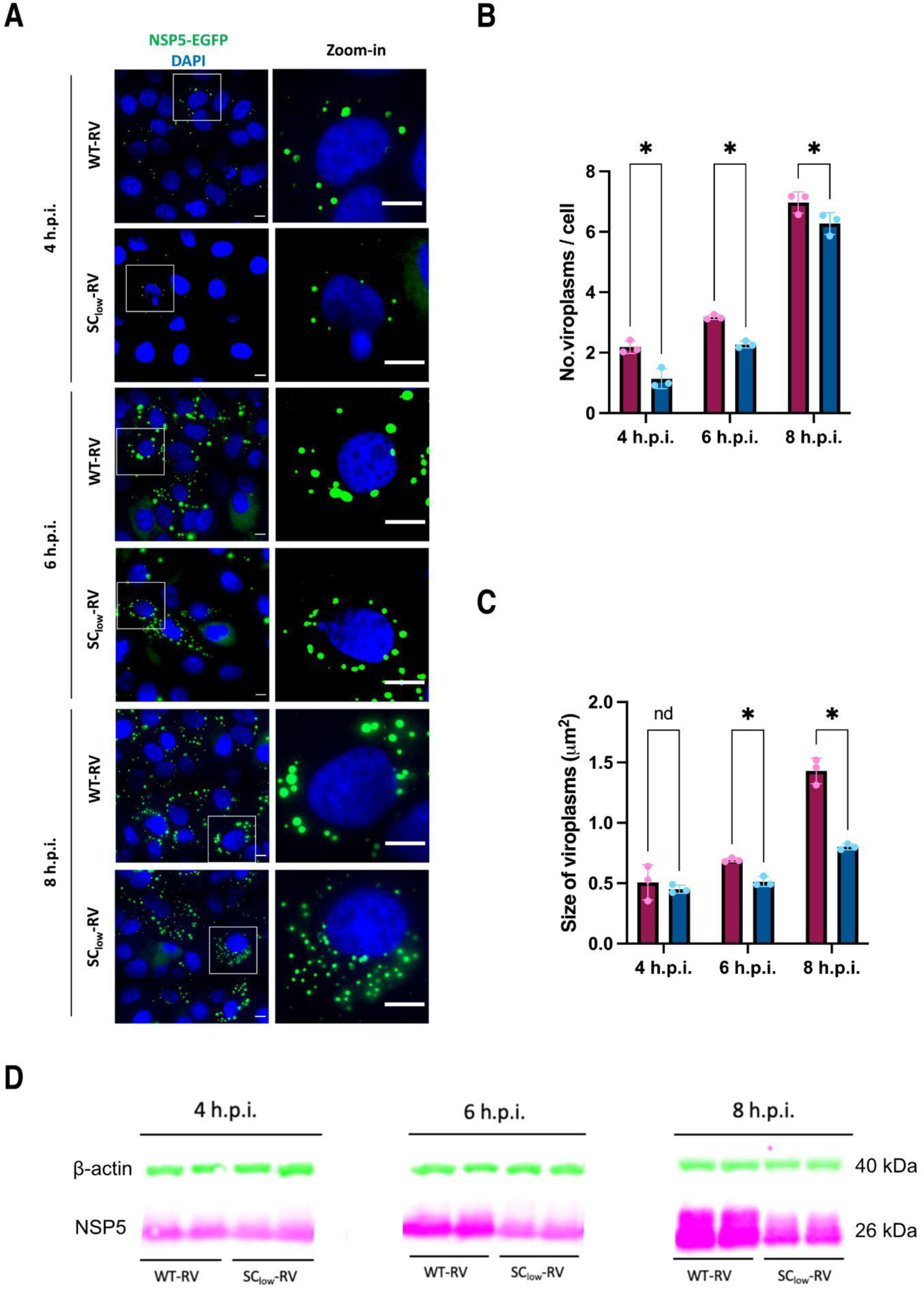
Delayed viroplasm formation and impaired NSP5 hyperphosphorylation in cells infected with SC_low_-RV. **A.** Wide-field fluorescence microscopy of MA104-NSP5-eGFP cells infected with either SA11-RV (WT-RV) or SC_low_-RV at MOI of 20. NSP5-eGFP–labelled viroplasms (green) and nuclei (DAPI, blue) are shown at 4, 6, and 8 hours post-infection (h.p.i.). Insets show enlarged regions of interest. Scale bar, 10 µm. **B,C.** Quantification of the number (**B**) and size (**C**) of viroplasms formed in cells infected by SA11-RV (burgundy) or SC_low_-RV (dark blue). Data represent mean values from three independent experiments (N > 200 cells per condition). Error bars indicate standard error of the mean (SEM). Two-way ANOVA was used to compare SA11-RV and SC^low-RV at each time point: viroplasm number at 4 h.p.i. (*p = 0.0004), 6 h.p.i. (*p = 0.0012), 8 h.p.i. (*p = 0.0071); viroplasm sizes at 4 h.p.i. (ns = 0.9244), 6 h.p.i. (*p = 0.0278), 8 h.p.i. (*p = 0.0211). **D.** Immunoblot analysis of NSP5 extracted from MA104 cells infected with SA11-RV or SC_low_-RV at 4, 6 and 8 h.p.i. (MOI = 20). Note the absence of the strongly hyperphosphorylated 35 kDa NSP5 isoform in SSC_low_-RV–infected cells at later time points.

**Expanded view Figure 4.**
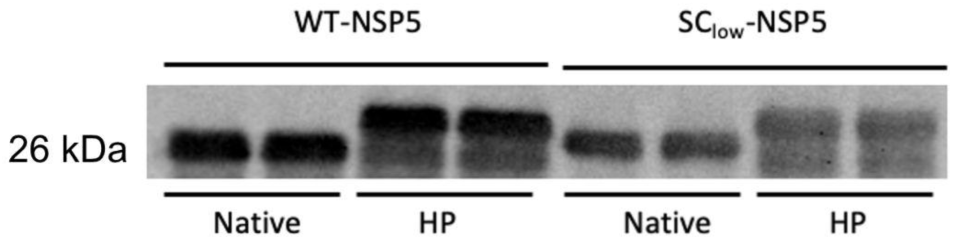
Validation of anti-NSP5 antibodies detecting both WT-NSP5 and SC_low_, as well as their phosphomimetics. Quantitative Western Blot analysis of recombinantly produced NSP5 (SA11 and SC_low_) and their corresponding hyperphosphorylation mimetics. Four distinct proteins (NSP5, SC_low_, NSP5 HP, and SC_low_ HP) were recombinantly produced and purified, as described in Methods. Equal amounts of each protein (400 ng per lane) were loaded in technical duplicates, resolved by SDS–PAGE, and analysed by Western blot using anti-NSP5 antibodies (Geiger et al., 2021).

Given that hyperphosphorylation of NSP5 increases progressively during infection (Papa et al., 2019), we next investigated whether SC_low_ -RV SC^low-RV undergoes a similar phosphorylation trajectory. From 6 h.p.i. onward, a notable difference in NSP5 hyperphosphorylation emerged between the two strains. This was especially pronounced at 8 h.p.i., when the most highly phosphorylated isoform of NSP5 (migrating at ∼35 kDa) was absent in SC_low_-RV-infected cells (**Fig 4D**). These findings suggest that the delayed and reduced viroplasm formation observed in SC_low_-RV-infected cells may be linked to impaired NSP5 hyperphosphorylation.

### Phosphorylation Restores Phase Separation and Viroplasm Formation by a Low-LLPS NSP5 Variant

As viroplasms represent complex condensates containing RNAs and additional viral proteins, we investigated phase-separation properties of SC_low_ using a minimal *in vitro* system containing NSP2 and SC_low_. Unlike its wild type counterpart, SC_low_ failed to form condensates upon mixing with NSP2 (**Fig 5**), in agreement with DeePhase predictions. Given that NSP5 undergoes progressive phosphorylation during RVA infection (Papa et al., 2019), and that we observed differences between WT-NSP5 and SC_low_ NSP5 phosphorylation patterns, we next asked whether phosphorylation could restore the ability of SC_low_ to phase separate *in vitro*. We introduced eight Ser-to-Asp mutations into both the WT-NSP5 (strain SA11) and the SC_low_ sequence (**Fig 2C**), targeting residues previously identified as phosphorylation sites in hyperphosphorylated NSP5 (Sotelo et al., 2010). In addition, we generated recombinant NSP5 from the bovine RVA strain RF (DeePhase score of 0.61), previously shown to undergo phase separation both in vitro and in cells (Geiger et al., 2021). These hyperphosphomimetic variants - derived from SA11, RF, or SC_low_ are referred to as NSP5 SA11 HP, NSP5 RF HP and SC_low_ HP, respectively.

**Figure 5.**
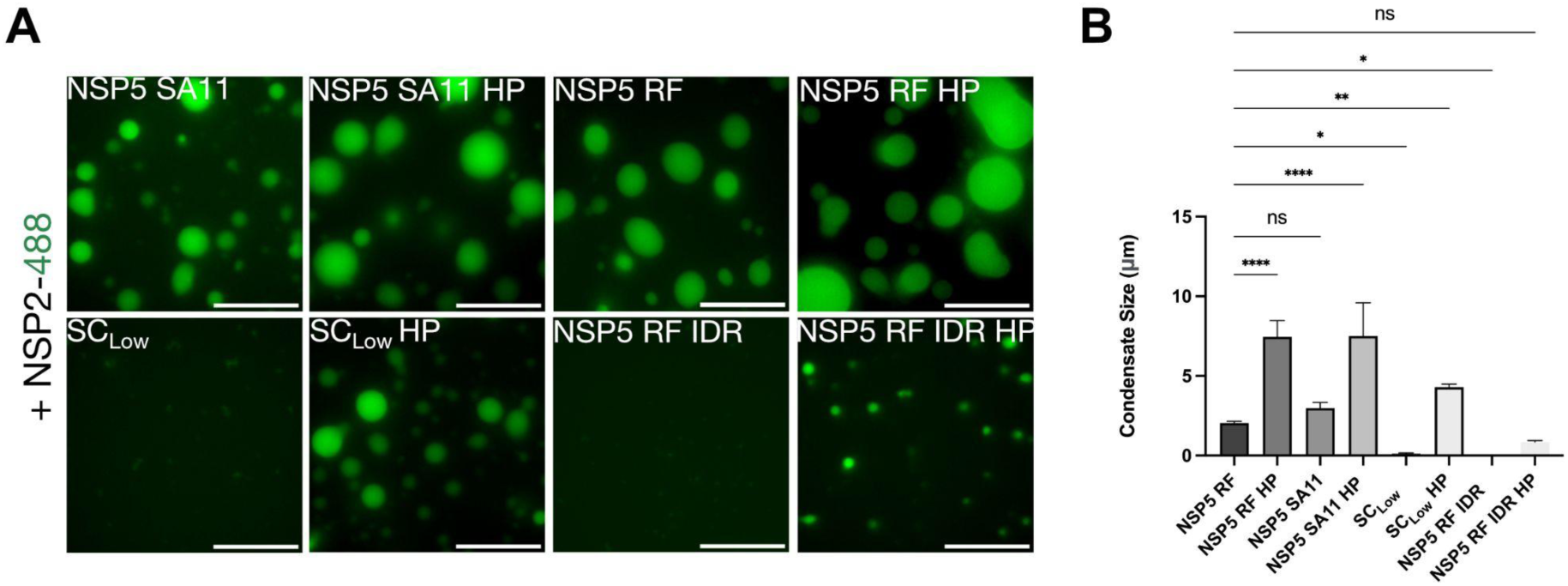
Hyperphosphorylation of SClow restores its ability to phase separate *in vitro*. **A.** *In vitro* phase separation assays of NSP5 variants in the presence of NSP2. Recombinant Atto-488-labelled NSP2 (25 µM) was mixed with equimolar amounts (25 µM) of unlabelled NSP5 variants, including wild-type SA11 (NSP5 SA11), its hyperphosphomimetic (NSP5 SA11 HP), bovine strain RF (NSP5 RF), NSP5 RF HP, SC_low_, SC_low_ HP, and the intrinsically disordered region-only construct of RF NSP5 (NSP5 RF IDR and IDR HP), as described in Methods. Fluorescence microscopy images show representative fields of view. Scale bar, 10 µm. **B.** Quantification of average condensate size for each NSP5 variant mixed with NSP2. Condensates were imaged within a 500 µm × 800 µm region of interest across three biological replicates (N = 3). Data were analysed by one-way ANOVA with Bonferroni and Sidak multiple comparisons test (α = 0.05). ****P < 0.0001.

Unlike SC_low_, the SC_low_ HP variant formed condensates upon mixing with low micromolar concentrations of NSP2 (**Fig 5**), confirming that phase separation of SC_low_ is phosphorylation dependent.

We observed that condensates formed by all HP variants were significantly larger than those formed by their non-phosphorylated counterparts (**Fig 5B**). Moreover, both SA11 and RF HP variants formed condensates at lower concentrations (i.e., reduced saturation concentration, C_sat_) and were significantly larger than those formed by SC_low_ HP (**Fig 6**).

**Figure 6.**
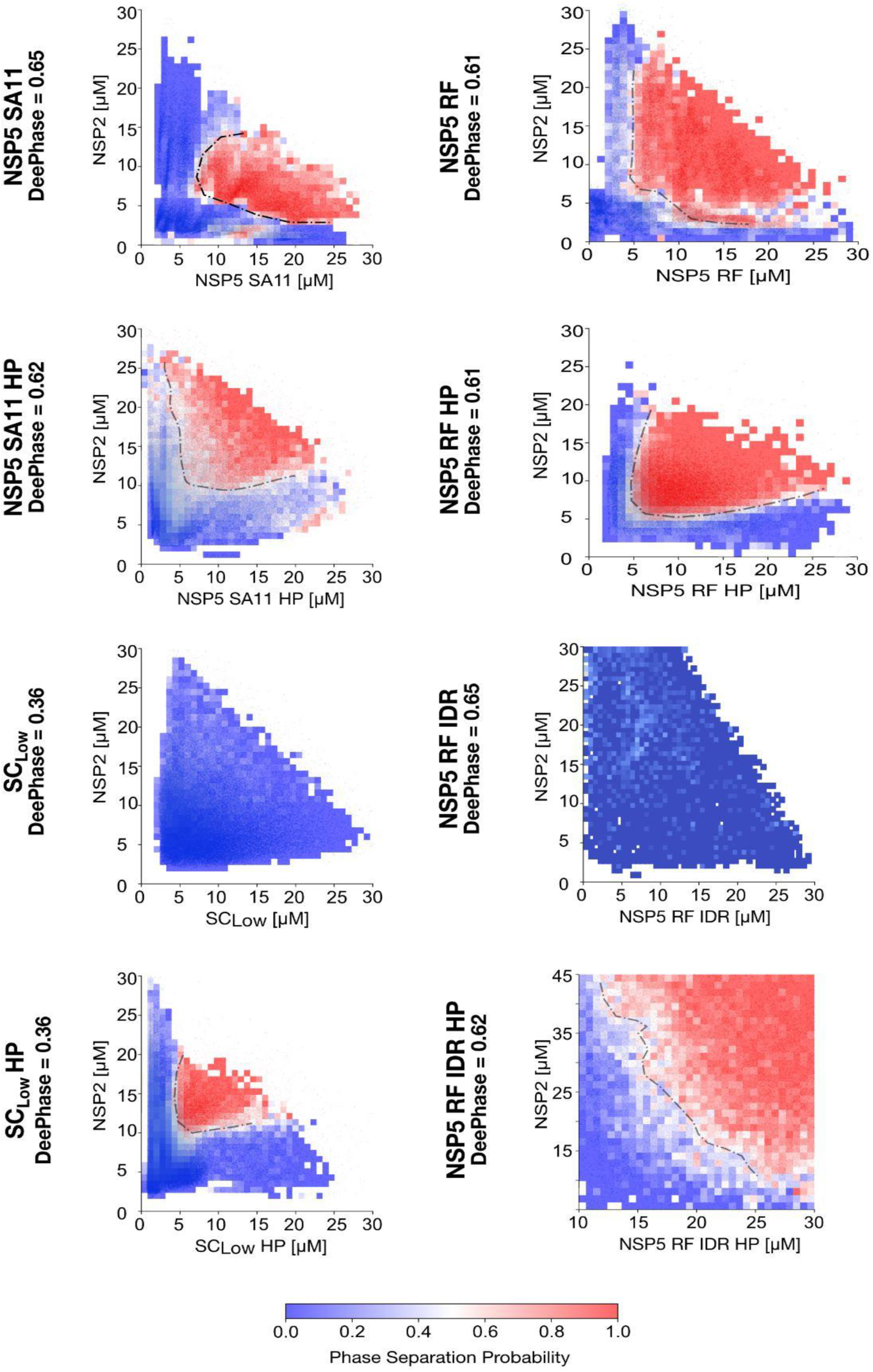
Phase behaviour of NSP5 variants in the presence of NSP2 reveals phosphorylation-dependent coacervation. PhaseScan-derived phase diagrams showing coacervation between NSP2 and different NSP5 variants. Combinatorial microfluidics was used to generate high-resolution phase diagrams for Atto-488-labelled NSP2 and Alexa-647-labelled NSP5 variants across a concentration matrix (2–25 µM for each protein), as detailed in Methods. LLPS probability heatmaps were reconstructed from the following number of droplets: NSP5 SA11 (n = 28,280), NSP5 SA11 HP (n = 87,053), SClow (n = 91,057), SClow HP (n = 101,659), NSP5 RF (n = 18,813), NSP5 RF HP (n = 17,449), NSP5 IDR (n = 36,511), and NSP5 IDR HP (n = 66,455). LLPS probability is colour-coded from low (blue) to high (red), with black dotted lines indicating inferred phase boundaries.

LLPS driven by homotypic interactions is characterised by a well-defined C_sat_, above which dense phases grow continuously as more protein is added. This behaviour is evident in the phase diagram of non-phosphorylated NSP5, which shows phase boundaries parallel to the concentration axes without re-entrant behaviour (**Fig 6)**. In contrast, LLPS driven by heterotypic interactions can be saturated by an excess of either component, leading to re-entrant behaviour, where condensates form and then dissolve as the stoichiometric ratio changes. This behaviour was observed in hyperphosphorylated NSP5 variants, including SC_low_ HP, which display curved phase boundaries (**Fig 6**).

To better understand the mechanistic basis of these differences, we examined NSP5 structural models generated by AlphaFold2 (Jumper et al., 2021) since no experimental structures of NSP5 are available. Both WT and SC_low_ monomers displayed similar features, notably a putative C-terminal alpha-helical region (CTR) and a largely disordered N-terminal region, which were also preserved in all HP variants (**Fig 7A**). As these monomeric structures did not reveal major differences between variants, and since oligomerisation of multivalent scaffold proteins can nucleate LLPS, e.g., by altering stoichiometry or affinity (Banani et al., 2017), we next assessed the oligomerisation capacity of the NSP5 variants using mass photometry. NSP5 SA11, RF, and their HP variants formed a range of higher-order oligomers, whereas SC_low,_showed only a single peak at 35 ± 5 kDa (**Fig 7B**). Remarkably, the SC_low_ HP variant formed a similar distribution of higher-order oligomers, confirming that phosphorylation is required for both oligomerisation and phase separation in this variant.

**Figure 7.**
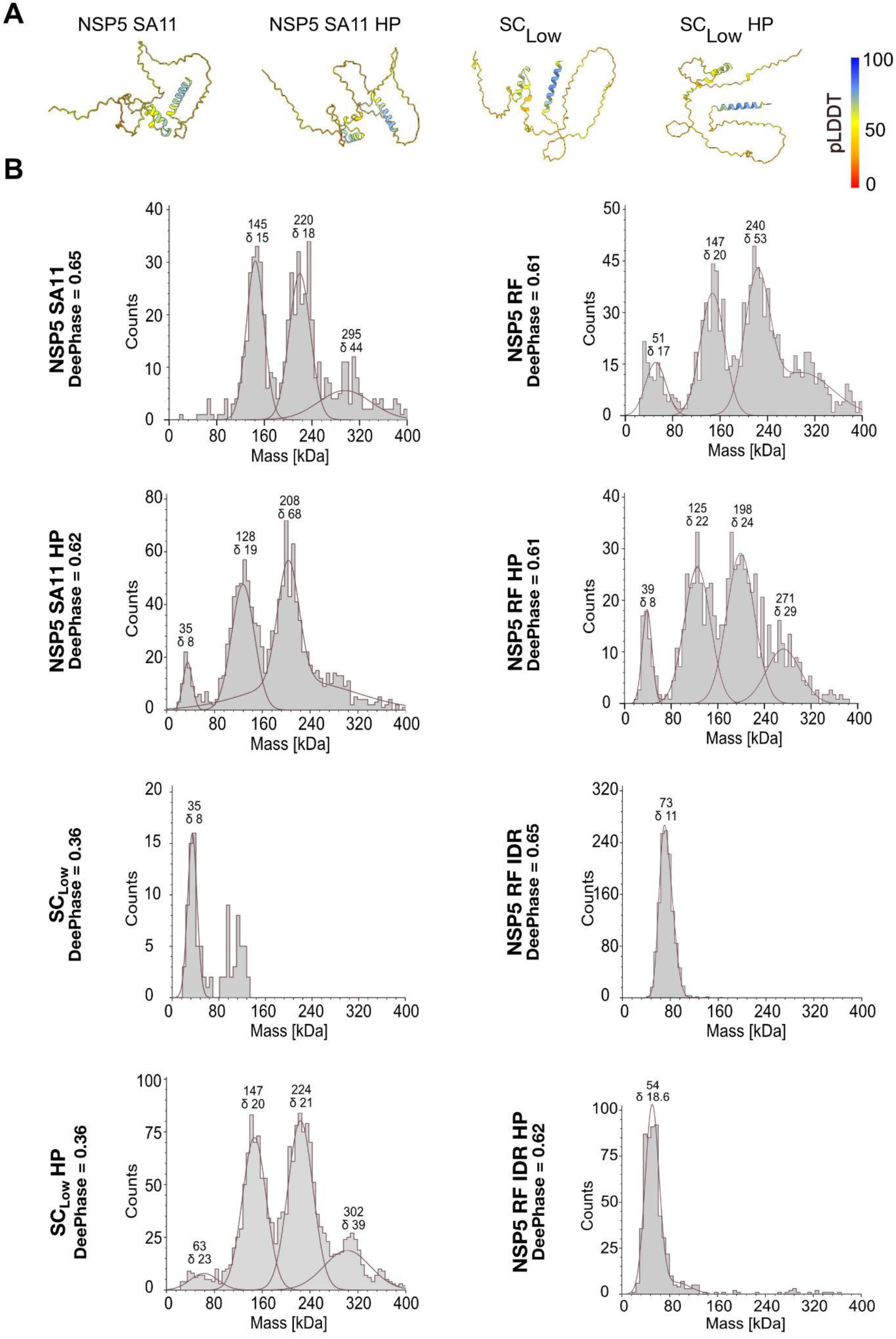
Hyperphosphorylation enables oligomerisation of SC_low_. **A.** AlphaFold2 structural models for NSP5 SA11, NSP5 SA11 HP, SC_low_ and SC_low_ HP. Models are coloured by per-residue confidence scores (pLDDT), with blue (pLDDT > 90) indicating high confidence and orange-red (pLDDT < 50) marking regions predicted to be intrinsically disordered. **B.** Oligomeric size distributions of NSP5 variants determined by mass photometry. Recombinant NSP5 SA11, NSP5 SA11 HP, SC_low_, SC_low_ HP, NSP5 RF, NSP5 RF HP, or NSP5 RF IDR were analysed at 100 nM each, as described in Methods. Representative molecular weight histograms are shown, with Gaussian fits highlighting major oligomeric species. Approximate molecular weights (in kDa) and corresponding standard deviations (σ) are indicated.

**Expanded View Figure 7.**
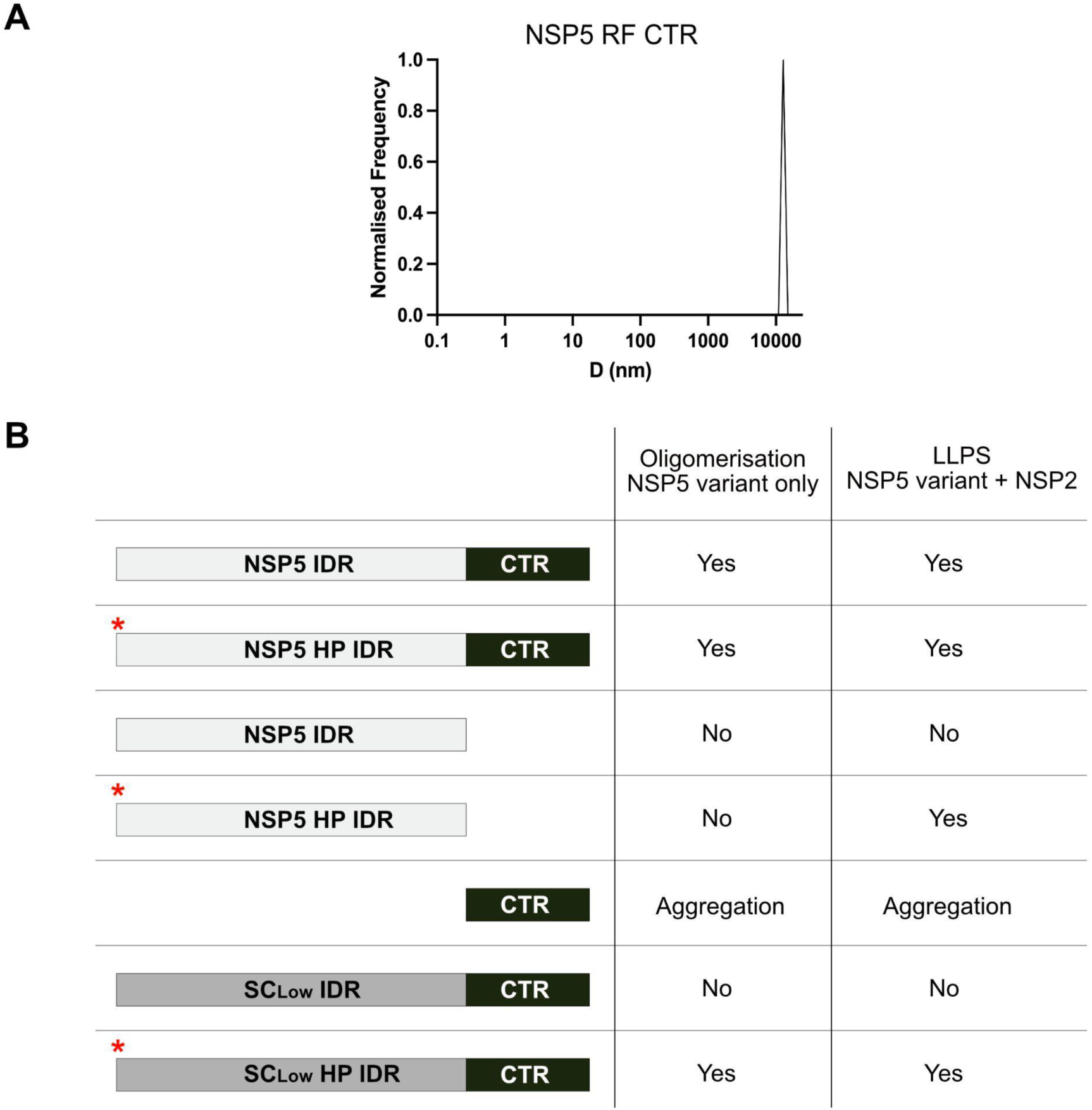
The C-terminal region (CTR) of NSP5 alone forms large aggregates. **A.** Hydrodynamic diameters (D, in nm) of 1 μM NSP5-RF CTR peptide measured in PBS (pH 7.4) using dynamic light scattering (DLS). **B.** Summary of the oligomeric states and liquid–liquid phase separation (LLPS) capacity of various NSP5 constructs used in this study. Schematic representations of the protein constructs are shown; phosphomimetic variants are indicated with a red asterisk.

Given that phosphorylation enables phase separation of SC_low_, we hypothesised that abrogation of phosphorylation would impair viroplasm formation and viral propagation in cells infected with the SC_low_-RV. To test this, we introduced the S67A mutation, previously shown to abrogate NSP5 hyperphosphorylation (Papa et al., 2019), into the SC_low_ background to generate the SC_low_ S67A recombinant RVA (**Fig 8A**). While the SA11 S67A mutant was able to replicate and form viroplasms in MA104 cells, the SC_low_ S67A virus could only propagate in MA104-NSP5 cells expressing SA11 NSP5 *in trans*, and failed to form visible plaques (**Fig. 8B**). Although both SA11 S67A and SC_low_ S67A viruses produced viroplasms at 24 h.p.i., only the former yielded visible condensates; no viroplasms were detected following SClow S67A infection (**Fig 8C**). Nonetheless, RV transcripts accumulated in MA104 and MA104-NSP5 cells infected with SC_low_ S67A-RV, indicating that viral transcription occurred in both contexts (**Fig 8D**).

**Figure 8.**
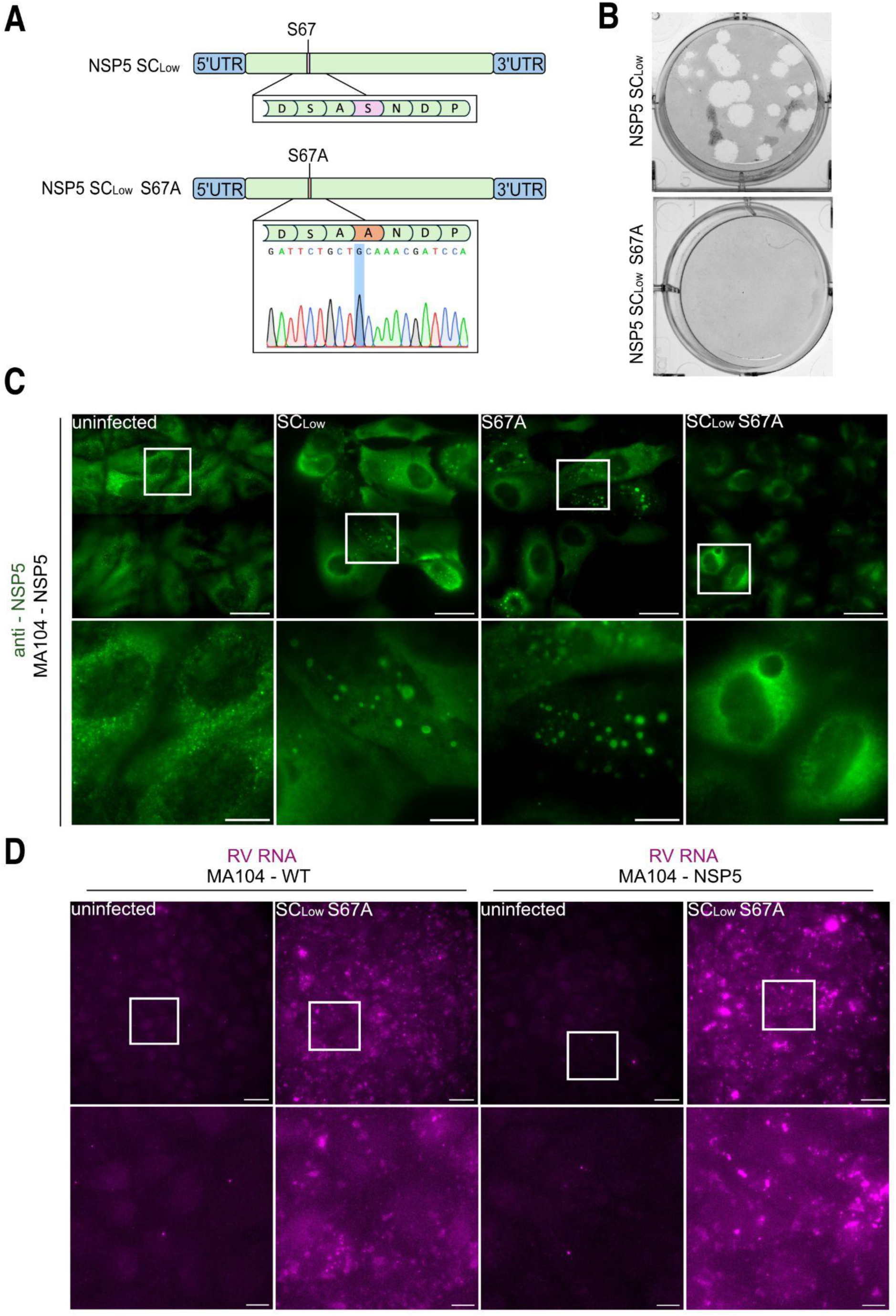
The SC_low_ S67A-RV mutant fails to form viroplasms in infected cells despite active viral transcription. **A.** Schematic representation of the S67A mutation introduced into gene segment 11 of SC_low_, with corresponding Sanger sequencing chromatograms confirming the mutation. **B.** Representative plaque assays in MA104-NSP5 cells infected with SC_low_-RV or SC_low_ S67A-RV. **C.** Wide-field fluorescence microscopy of NSP5-eGFP-tagged viroplasms (green) in cells infected with SC_low_-RV, SA11 S67A-RV or SC_low_ S67A-RV at an MOI of 10. Images were acquired at 24 h.p.i. Scale bars: overview = 30 µm; zoomed regions = 10 µm. **D.** RNA FISH detection of viral transcripts (gene segment 1) in MA104 WT and MA104-NSP5 cells infected with SC_low_ S67A-RV, using Atto647N-labelled probes. Viral RNA accumulates in both cell types, indicating transcriptional activity. Scale bars: overview = 30 µm; zoomed regions = 10 µm.

Together, these findings support the conclusion that phosphorylation is critical for SC_low_ NSP5 to undergo LLPS, form viroplasms, and support efficient rotavirus replication.

### Phosphorylation tunes NSP5 binding modes in condensates

Phosphorylation events in NSP5 are confined to the intrinsically disordered region (IDR) (**Fig 2C**) (Sotelo et al., 2010). While the NSP5 IDR has been primarily implicated in mediating NSP2–NSP5 interactions (Eichwald et al., 2004b; Fabbretti et al., 1999) prior studies, including ours, have shown that the CTR is essential for NSP5 self-association (Geiger et al., 2021; Martin et al., 2011). However, the absence of homo-oligomerisation in SC_low_ suggests that the IDR may also contribute to NSP5 self-association.

To investigate this further, we analysed the CTR peptide alone using dynamic light scattering, which revealed the formation of large aggregates (**Expanded View Fig 7A**). This finding suggests that the IDR normally inhibits uncontrolled CTR aggregation, implying that both regions act cooperatively in oligomerisation and phase separation. Consistent with this, deletion of the CTR (yielding the IDR-only construct) abolished homo-oligomerisation (**Fig 7B**) and eliminated NSP2-driven phase separation (**Fig 5**) (Geiger et al., 2021). To dissect the contributions of each domain to condensate formation, we titrated increasing concentrations of the isolated IDR or CTR into preformed NSP5/NSP2 condensates. As expected, the CTR strongly disrupted condensate nucleation, reflecting its aggregation-prone nature. The IDR also impaired condensate formation, though requiring a ∼3-fold higher concentration. Notably, replacing NSP5 with its phosphomimetic variant (NSP5 HP) conferred greater tolerance to the CTR (**Fig 9**), suggesting that phosphorylation of the IDR modulates homotypic interactions within condensates. In all cases, condensate size decreased when full-length NSP5 concentration was reduced, indicating that both termini are required for efficient condensate growth (**Fig 9**).

**Figure 9.**
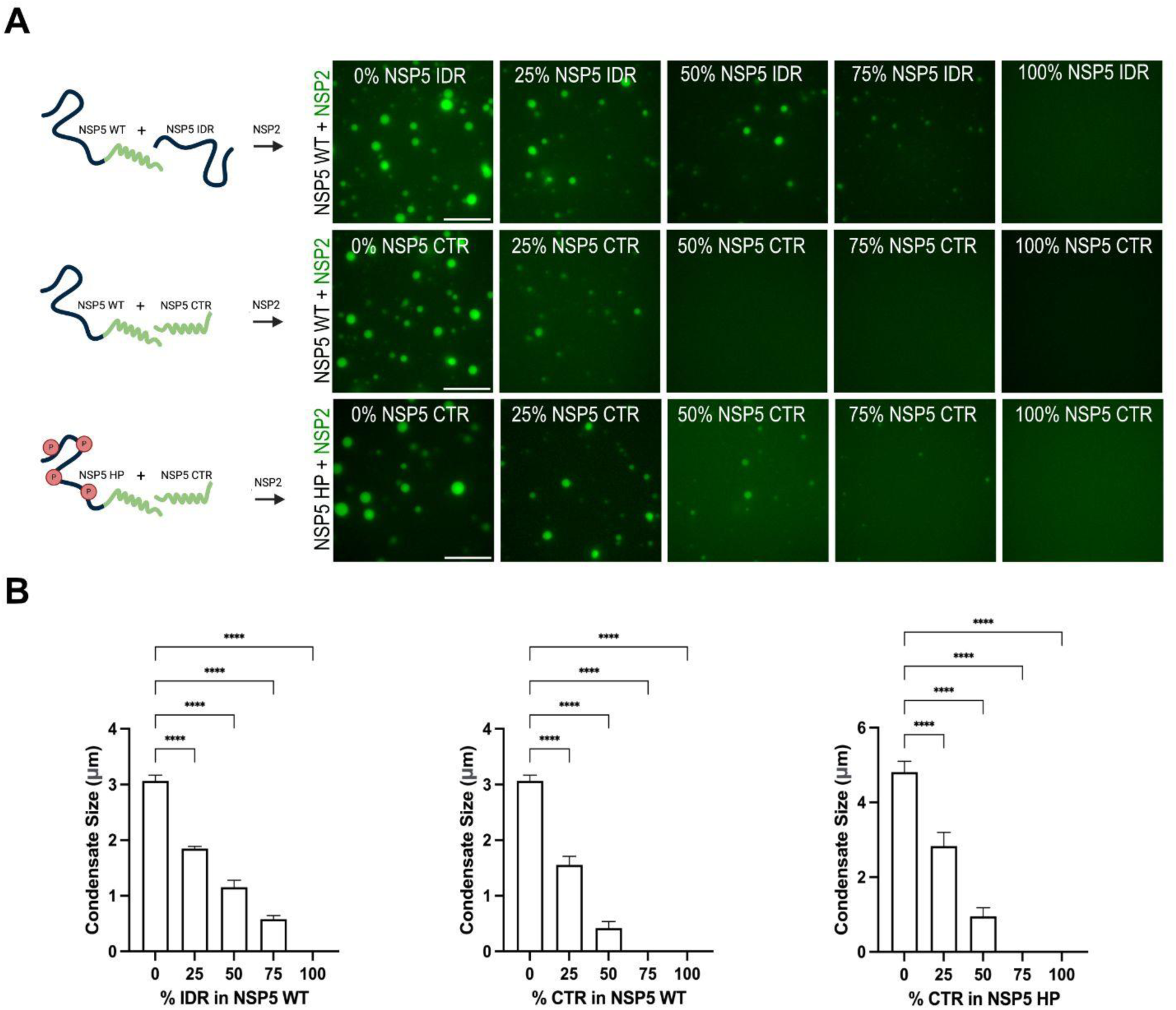
Both the NSP5 CTR and intrinsically disordered region (IDR) are critical for mediating LLPS. **A.** Experimental setup to assess the individual contributions of the NSP5 IDR and CTR to phase separation. Wild-type NSP5 or its phosphomimetic variant (NSP5 HP) was mixed with increasing ratios of either the isolated IDR or CTR, followed by addition of NSP2 to induce LLPS. Schematics illustrate NSP5 domains: IDR (black), CTR (green), and phosphomimetic sites (red ’P’). Phase-separated condensates were imaged by wide-field microscopy following mixing of Atto-488-labelled NSP2 (25 μM) with total NSP5 (25 μM) at the indicated full-length:CTR or full-length:IDR ratios. Scale bar, 10 μm. **B.** Quantification of condensate area (μm²) formed by NSP2 with wild-type or HP NSP5 in the presence or absence of IDR or CTR fragments. Nine regions of interest (50 µm × 80 µm) were analysed per condition, as described in Methods. Statistical analysis was performed using one-way ANOVA with Dunnett’s multiple comparisons test (α = 0.05): **P = 0.0082, ***P = 0.0003, ****P < 0.0001.

**Expanded View Figure 9.**
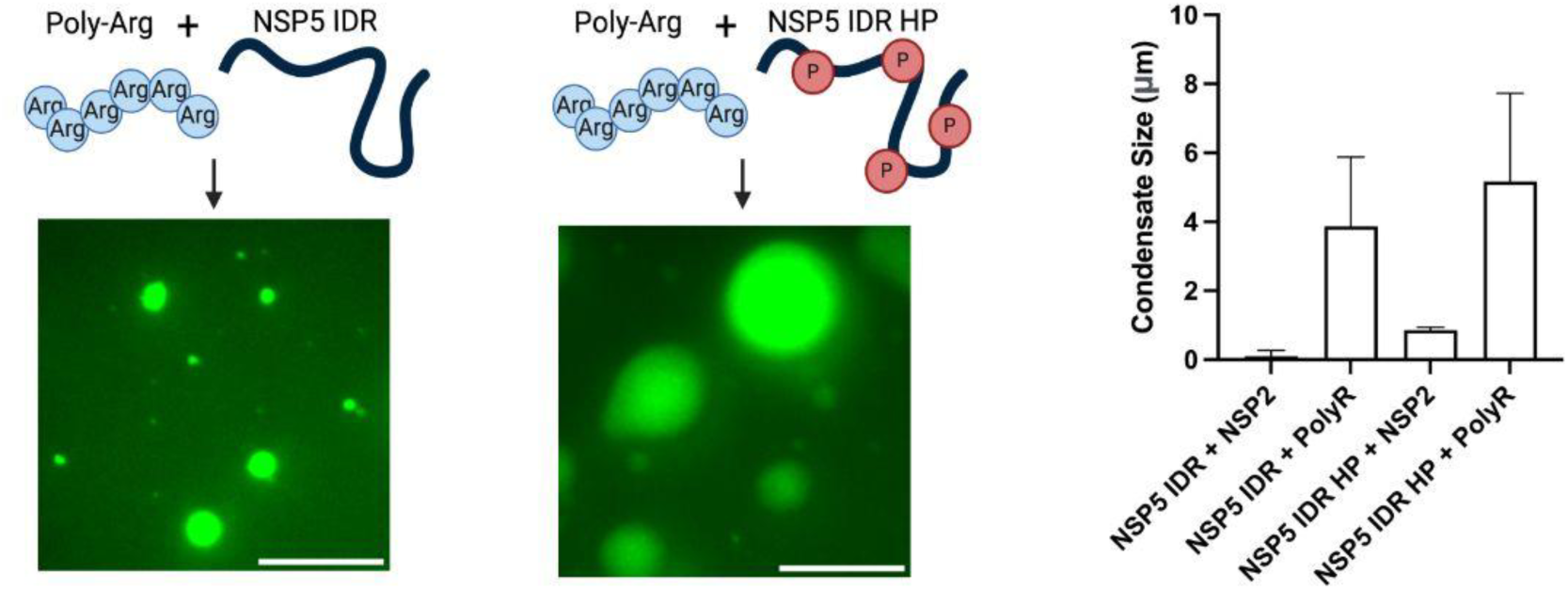
Both phosphorylated and non-phosphorylated NSP5 IDRs undergo phase separation with poly-arginine. **A.** Phase separation of NSP5 IDR and its phosphomimetic variant (IDR HP) in the presence of poly-L-arginine. DyLight-488-labelled IDR or IDR HP (25 μM) was mixed with 5 μM poly-L-arginine (schematically illustrated), and resulting condensates were imaged by wide-field microscopy. Phosphomimetic sites are indicated in red (‘P’). Scale bar, 10 μm. **B.** Quantification of condensate area (μm²) formed by NSP5 IDR or IDR HP with poly-L-arginine. Nine regions of interest (50 µm × 80 µm each) were analysed as described in Methods. For comparison, condensate sizes formed by NSP2 with NSP5 IDR or IDR HP (as shown in Figure 9) are also included.

Given that SC_low_ fails to phase separate unless phosphorylated and resembles the IDR in its oligomeric behaviour, we hypothesised that phosphorylation of the IDR also regulates its interaction with NSP2. To test this, we introduced phosphomimetic mutations into the RF-derived NSP5 IDR (IDR HP). Like its unmodified counterpart, IDR HP failed to form higher-order oligomers (**Fig 7B**). However, in the presence of NSP2, IDR HP was able to phase separate, albeit forming smaller condensates and requiring ∼4-fold higher concentrations than WT NSP5 (**Fig 6**). These data indicate that IDR phosphorylation is required for co-condensation with NSP2, but that the CTR remains essential for efficient condensation, as reflected in the elevated C_sat_ of IDR HP.

Since non-phosphorylated IDR does not interact with NSP2, we hypothesised that phase separation of IDR HP is largely driven by electrostatic interactions with the positively charged NSP2. To test this, we examined whether poly-arginine could induce condensation of the IDRs via cation–π interactions (Geiger et al., 2021). While NSP2 failed to induce condensation with the non-phosphorylated IDR, poly-arginine efficiently promoted condensate formation with both IDR variants (**Expanded View Fig 9**). These results indicate distinct mechanisms of condensation for phosphorylated versus unphosphorylated NSP5 in the presence of NSP2.

To map phosphorylation-dependent binding interfaces, we performed hydrogen– deuterium exchange mass spectrometry (HDX-MS) on WT NSP5 and its phosphomimetic variant in the presence of NSP2 (**Fig 10**). In unmodified NSP5, NSP2 binding led to significant protection of both the CTR (residues 167–191) and an IDR region (residues 26–47), in agreement with previous reports of NSP2–NSP5 interaction sites (Eichwald et al., 2004b; Lee et al., 2024). In contrast, the NSP5 HP variant showed no protection at these sites. Instead, a different region within the IDR (residues 53–73) became protected, confirming that NSP5 uses distinct binding interfaces depending on its phosphorylation state.

**Figure 10.**
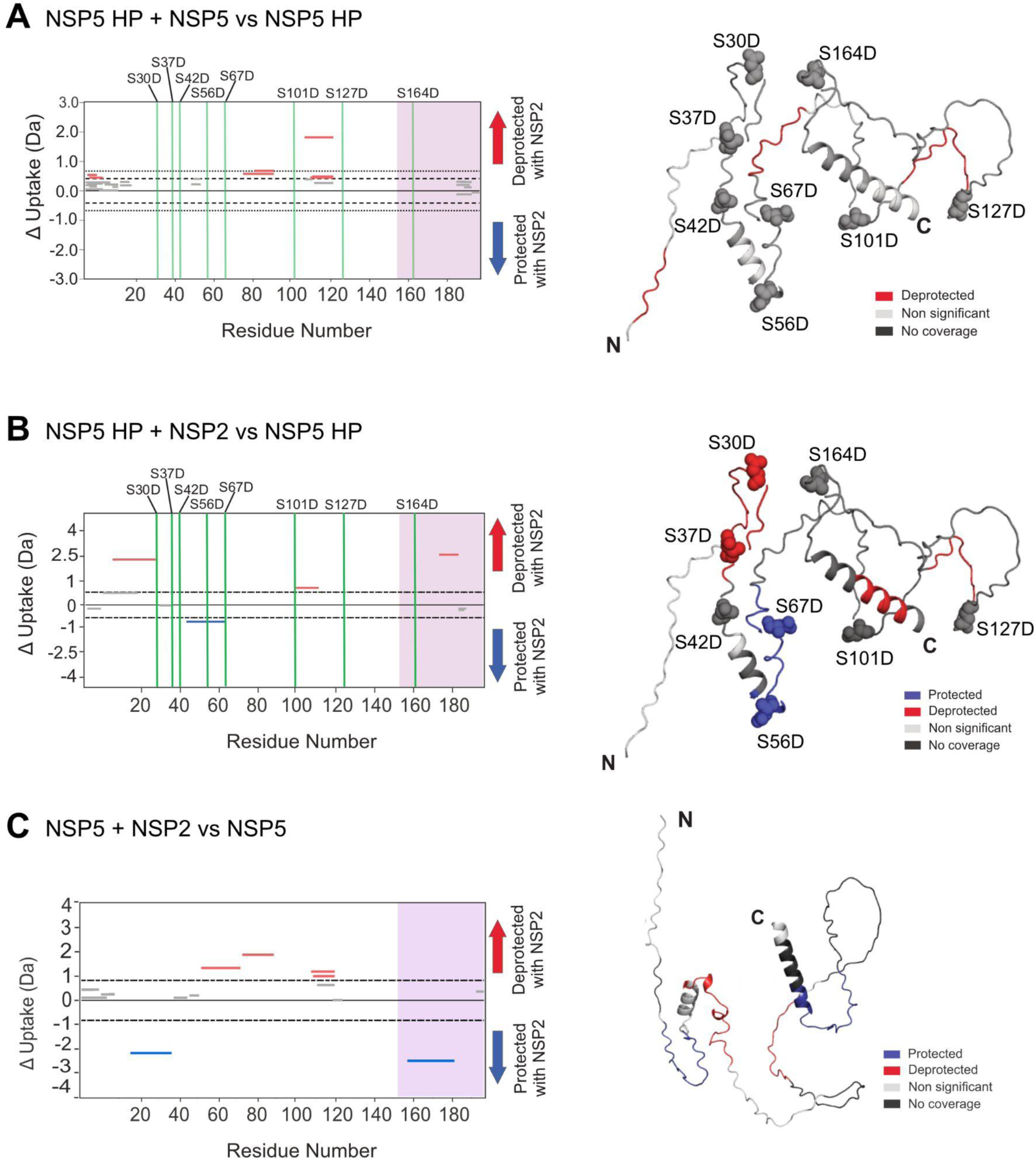
Distinct NSP5–NSP2 interaction interfaces revealed by HDX-MS upon hyperphosphorylation of NSP5. **A–C.** Hydrogen–deuterium exchange mass spectrometry (HDX-MS) analysis of wild-type (WT) and hyperphosphorylation-mimetic (HP) NSP5 in the absence and presence of NSP2. Cumulative Woods plots show peptides of NSP5 HP (**A, B**) and WT NSP5 (**A, C**) that exhibit significant changes in deuterium uptake upon incubation with deuterium alone or with NSP2, analysed using Deuteros with hybrid significance testing (p < 0.02; (Lau et al., 2019)). Regions showing increased protection (blue) or deprotection (red) are indicated. The C-terminal region (CTR) is marked by a purple box. **D.** AlphaFold2 model of monomeric NSP5 showing mapped HDX-MS data: protected regions (blue), deprotected regions (red), peptides with no significant change (light grey), and regions with no coverage (dark grey). Ser-to-Asp phosphomimetic mutations are indicated.

## Discussion

Although many IDPs can spontaneously phase separate, others require post-translational modifications such as phosphorylation to facilitate condensation (Aumiller and Keating, 2016; Grams et al., 2024). Given the extensive sequence diversity across viral proteomes (Dyson, 2023; Mishra et al., 2020), we hypothesised that different viral variants might employ these mechanisms interchangeably depending on their amino acid composition of their IDPs.

We identified NSP5 sequences with markedly different condensate-forming abilities compared to the well-characterised rotavirus strain SA11. This allowed us to engineer a low LLPS propensity NSP5 variant, SC_low_, which retains conserved features of the SA11 sequence. SC_low_ showed a significantly reduced DeePhase score achieved through combinatorial amino acid substitutions derived from four naturally occurring NSP5 sequences with similarly low scores.

DeePhase accurately predicted LLPS potential: both SA11 NSP5 and SC_low_ behaved as expected *in vitro.* Paradoxically, SC_low_ fully supported viral replication, forming viroplasms in infected cells. While DeePhase and similar approaches are trained on proteins with known propensity to phase separate (Chu et al., 2022; Hatos et al., 2023; Saar et al., 2021), they do not reveal specific features or residues driving LLPS. Our combined use of in silico prediction and mutational analysis revealed that DeePhase is sensitive primarily to sequence differences within IDRs, but not to contributions from structured regions, which may still be critical for phase separation. Notably, phosphomimetic substitution of serines previously identified as phosphorylated in infected cells, restored the ability of SC_low_ to nucleate LLPS similarly to the wild-type protein.

Previous studies identified the helical C-terminal region (CTR) of NSP5 as the key interface mediating its homo-oligomerisation (Martin et al., 2011; Sen et al., 2007; Torres-Vega et al., 2000). While, coiled-coil interactions can be sufficient to drive phase separation in other systems (Ramirez et al., 2023; Ramšak et al., 2023), the inability of SC_low_ to undergo LLPS despite an intact CTR demonstrates that both the IDR and the CTR are required for phase separation. Indeed, deletion of the CTR abrogates high molecular weight NSP5 species, while removal of the IDR results in aggregation of the CTR, indicating that both regions contribute to NSP5 oligomerisation (**Fig 11A**).

**Figure 11.**
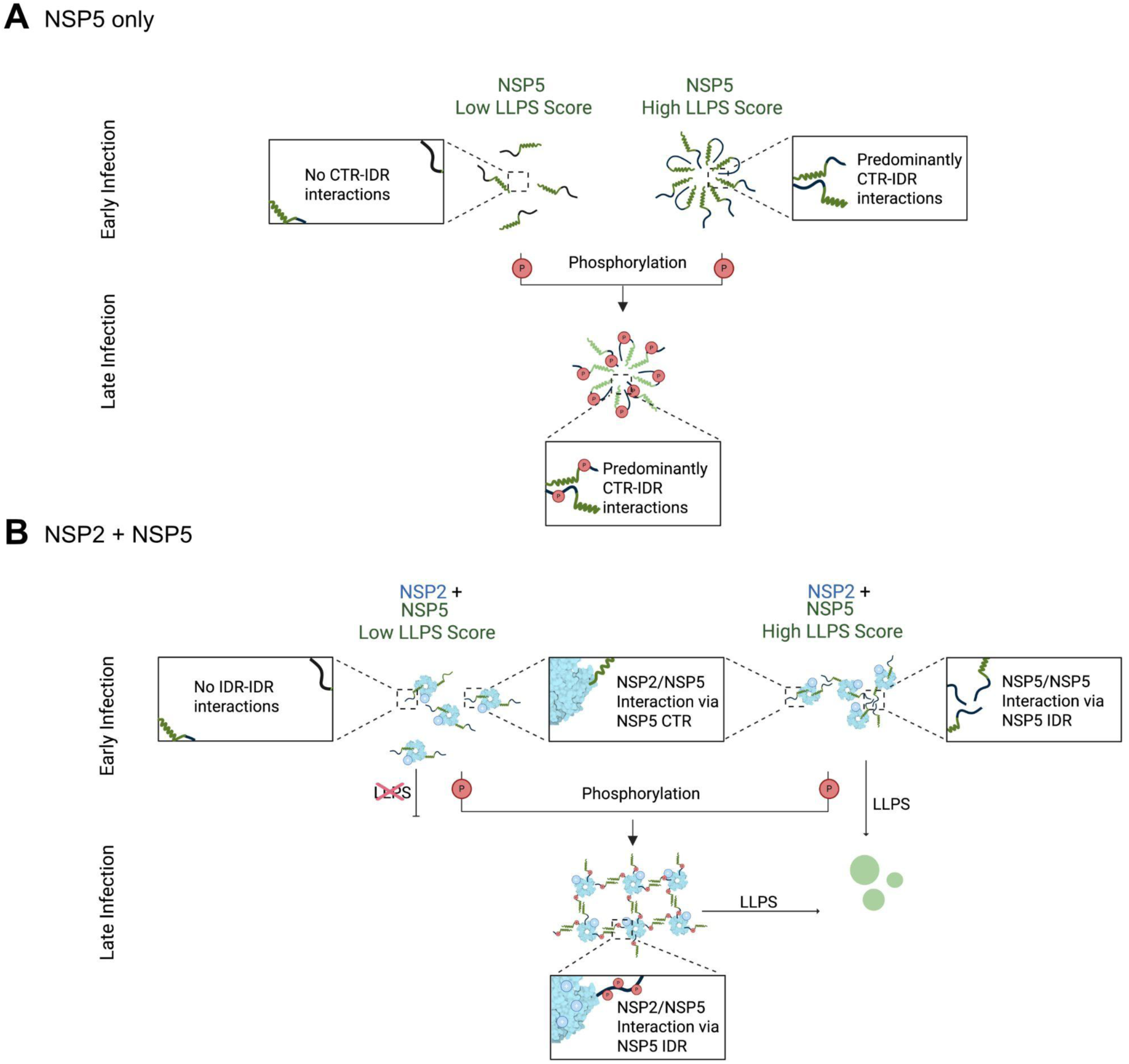
Phosphorylation-induced transition from non-saturating to saturating interactions in NSP5-NSP2 condensates. **A.** Homotypic interactions of NSP5. NSP5 molecules with low LLPS scores (black IDRs and green CTRs) are unable to self-associate effectively. In contrast, NSP5 variants with high LLPS scores or those containing phosphorylated IDRs can form condensates, with LLPS favoured by interactions between the IDR and CTR. Phosphorylation of the IDR enhances these IDR–CTR interactions, facilitating homo-oligomerisation and condensate formation **B.** Heterotypic interactions between NSP5 and NSP2. At early stages of infection, positively charged NSP2 octamers (light blue doughnuts) bind NSP5 via stoichiometric interactions with its CTR. In NSP5 variants with high LLPS propensity, the flexible, unstructured IDRs form a non-stoichiometric interaction network that drives phase separation. In contrast, low-LLPS NSP5 variants can still bind NSP2 via the CTR but fail to phase separate due to insufficient IDR–IDR interactions. During late infection, phosphorylation of NSP5 increases the negative charge of the IDR, promoting interactions with positively charged regions of NSP2. These phosphorylation-dependent interactions enable even low-LLPS NSP5 variants to engage in saturable binding and phase separation, facilitating condensate assembly.

Importantly, oligomerisation of NSP5 appears to be dispensable for NSP5/NSP2 co-condensation, as the IDR HP alone can phase separate with NSP2. However, oligomerisation significantly reduces the saturation concentration (Csat), suggesting that interactions promoting NSP5 self-assembly are distinct from those driving its co-condensation with NSP2 (Choi et al., 2020). This implies that sequence features within the IDR not only prevent CTR aggregation but also modulate the cooperative assembly of NSP5 oligomers, and NSP5 condensates.

We propose a model in which unphosphorylated NSP5 binds NSP2 via the CTR, forming a hub–driver architecture (Galagedera et al., 2023), where the remaining free IDRs mediate phase separation through non-stoichiometric interactions (**Fig 11**). This is consistent with our observation that the non-phosphorylated IDR alone cannot bind NSP2. Phosphorylation introduces negative charges predominantly within the IDR, which may either weaken homotypic IDR–IDR interactions or enhance binding to a distinct region of NSP2, driven by electrostatic attraction to its positively charged domains. As a result, the mechanism of phase separation may shift from weak, non-saturating multivalent interactions to specific, saturable ones (**Fig 11B**), representing an allosteric switch in NSP5’s interaction mode. Upon phosphorylation, the IDR binds NSP2, and homotypic interactions occur primarily through the CTR. Unlike IDR-mediated interactions, CTR-based oligomerisation is saturable and can be competitively inhibited by excess NSP5, leading to condensate dissolution. These findings align with theoretical models proposing that condensate function is governed by a balance of weak, competing interactions (Bhandari et al., 2021; Schmit et al., 2021), which are often elusive to conventional biophysical methods. Our data highlight how networks of weak, competing interactions within liquid-like biomolecular condensates can be dynamically regulated by phosphorylation.

We propose that phosphorylation-dependent switching between condensation mechanisms may be essential for different stages of the rotavirus life cycle, enabling viroplasms to transition from the initial recruitment of replication components to later assembly steps.

Although eight serines in NSP5 have been identified as phosphorylated (Sotelo et al., 2010), the minimum number required to trigger an allosteric switch remains unknown. Multi-site phosphorylation of the IDP Sic1 serves as a paradigm for a dose-dependent threshold response, as Sic1 possesses 9 distinct phosphosites, and phosphorylation of at least six of these is required for its interaction with its binding partner (Borg et al., 2007). Similarly, multiple phosphates reside within an IDR of ETS-1 transcription factor that functions as an allosteric effector of autoinhibition. Graded phosphorylation of ETS-1 serves as a rheostat to fine-tune DNA binding (Pufall et al., 2005), providing another example of an elegant control of phosphorylation-driven allosteric regulation in IDRs, that can dynamically tune the strength of such interactions.

Hyperphosphorylated NSP5 gradually accumulates during infection, remaining undetectable during early stages of infection (Blackhall et al., 1998; Campagna et al., 2007; Criglar et al., 2018; Eichwald et al., 2004a). Such phosphoregulation may be required for the advanced stages of viral replication as the amount of NSP5 and NSP2, as well as other viral components required for virus assembly is high. In the SA11 strain, which has high intrinsic LLPS propensity, NSP5 phosphorylation is not required for in vitro condensate formation or early-stage viroplasm formation (Papa et al., 2019). Conversely, SC_low_ lacking IDR-IDR interactions cannot initiate condensate formation unless phosphorylated. The S67A mutation, which blocks a key phosphorylation site, abolishes phase separation and viroplasm formation in the SC_low_ background. Thus, in NSP5 variants with low LLPS propensity, phosphorylation serves as a prerequisite for condensate nucleation and viral replication. Despite sequence divergence, phosphorylation enables these variants to form condensates, underscoring a conserved mechanism across rotavirus strains.

We propose that hyperphosphorylation of NSP5 serves dual roles depending on strain context. In low-LLPS variants, it enables LLPS nucleation; in high-LLPS strains, it likely alters NSP5’s interaction modes with other viroplasmic components, e.g., VP1, VP3, viral RNAs, and the inner core protein VP2.

Viral proteomes are rich in IDRs, offering evolutionary flexibility to adjust LLPS potential through the gain or loss of phosphorylation sites. This may help compensate for mutations that weaken condensate formation, while also facilitating the use of host-specific kinases. The ability of IDPs to switch interaction modes upon phosphorylation enables access to diverse functional states during replication and assembly. These properties make viral IDPs promising antiviral targets, particularly through the development of allosteric inhibitors. Indeed, such an approach has successfully targeted the IDR of a protein tyrosine phosphatase (Krishnan et al., 2014). Future studies into how viruses exploit IDR-driven condensation will offer new insights into viral evolution, host adaptation, and the regulatory logic of biomolecular condensates.

## Methods

### NSP5 Sequences

A dataset comprising 89,417 *Rotavirus A* protein sequences was obtained from the NCBI Virus database (Taxonomy ID: 28875). FASTA headers were reformatted to include relevant metadata (Accession, GenBank Title, Species, Length, Segment, and Protein designation), which facilitated filtering for full-length NSP5 sequences. A custom Python script (described below) was used to exclude entries marked as “truncated” or “partial,” as well as sequences containing ambiguous amino acid codes: “B” (aspartic acid or asparagine), “Z” (glutamic acid or glutamine), “J” (leucine or isoleucine), and “X” (unspecified residues). Sequences annotated as “NSP5” or its variants were retained. After removing duplicates, a curated set of 451 unique full-length NSP5 sequences was obtained.

### LLPS Propensity Prediction Using DeePhase

Liquid–liquid phase separation (LLPS) propensities of NSP5 sequences were assessed using DeePhase (Saar et al., 2021) and its associated predictive framework. DeePhase employs two independent models: the first relies on sequence-derived biophysical features, including hydrophobicity, Shannon entropy, sequence length, the proportion of polar, aromatic, and positively charged residues, as well as the content of low-complexity and intrinsically disordered regions. The second model is based on word2vec embeddings (Mikolov et al., 2013b, 2013a), a natural language processing algorithm adapted to capture sequence patterns. Both models were trained on the publicly available LLPSDB (Li et al., 2020) and PDB (Berman et al., 2000) datasets. Predictions were performed for a curated set of 451 unique full-length NSP5 sequences, including those from the SA11 and RF strains (NCBI accession IDs BAW94621.1 and AHF49898.1), 128 permutation variants, and the engineered low-propensity sequence, SClow. Each sequence was evaluated by both prediction models, and the final LLPS propensity score was calculated as the average of the biophysical and word2vec-based scores, following the authors’ recommendations (Table 1).

### Sequence Distance Calculation

Pairwise Levenshtein distances (Levenshtein, 1966) were computed between the SA11 NSP5 sequence (NCBI accession ID: BAW94621.1) and all NSP5 variants with a DeePhase propensity score below 0.5 (S1_low_, …, S7_low_; NCBI accession IDs QIN53369.1, QIN53336.1, ACC91692.1, BBB18699.1, QIJ58158.1, AGF92026.1, AKA40124.1). Distances were calculated using a custom Python script (v3.9.5) with the Python-Levenshtein package (v0.21.1) and plotted with matplotlib (v3.4.2). All custom scripts have been deposited and are available at https://github.com/desiro/SClow.

### Consensus Sequence Generation

Multiple sequence alignments were performed using MAFFT (v7.480) (Katoh and Standley, 2013a) with default parameters. The alignments were imported into Unipro UGENE (v42.0) (Golosova et al., 2014; Okonechnikov et al., 2012; Rose et al., 2019) to create consensus sequences. A consensus sequence was generated for (i) the four sequences from the SA11-like cluster (SC; s4_low_, s5_low_, s6_low,_ and s7_low_) and (ii) the subset of 78 SC128 permutation sequences with DeePhase liquid–liquid phase separation propensity scores below 0.5 (Saar et al., 2021).

### Phylogenetic Tree Construction

To construct the phylogenetic tree, all 451 unique rotavirus A NSP5 sequences retained after filtering, along with the 128 permutation sequences and the engineered SClow sequence (GenBank accession no. PP828582), were aligned using MAFFT (v7.480) (Katoh and Standley, 2013b) with default parameters. The resulting alignment was subjected to maximum likelihood phylogenetic analysis using PhyML 3.0 (v3.3.20200621) (Guindon et al., 2010) with the command-line options “--datatype aa --no_memory_check --leave_duplicates”. The phylogenetic tree was midpoint-rooted (Swofford et al., 1996) and visualised using the Interactive Tree of Life (iTOL v6) (Letunic and Bork, 2021).

### Genomic RNA Sequence Creation

The RNA sequence encoding SClow was generated using a custom Python script (v3.9.5; Python Software Foundation, 2001), with the SA11 NSP5 RNA sequence serving as the reference (see **Expanded View Fig 3A**). To ensure compatibility with the SA11 rotavirus genome, the algorithm preserved both 5′ and 3′ untranslated regions (UTRs) of gene segment 11, modifying only the coding region through synonymous codon substitutions. For codons where the encoded amino acid remained unchanged, the nucleotide sequence from the SA11 reference was retained. For codons corresponding to amino acid substitutions, synonymous codons were selected based on minimal evolutionary distance to the reference codon, using the K3ST model (Kimura, 1981) based on the generalised time-reversible model (Tavaré et al., 2000). The K3ST parameters were set to α = 1.4, β = 1.24, and γ = 1.36.

### Cells and Viruses

African green monkey foetal kidney MA104 cells (ATCC CRL-2378.1), which are susceptible to rotavirus (RV) infection (simian SA11 strain G3P[2]), were used for virus propagation. For imaging experiments, a derivative MA104 cell line stably expressing NSP5-eGFP (NSP5-eGFP-MA104) was used (Papa et al., 2020). Both wild-type MA104 and NSP5-eGFP-MA104 cells were cultured as monolayers in Dulbecco’s Modified Eagle Medium (DMEM, Gibco), supplemented with 10% fetal bovine serum (FBS; Sigma-Aldrich), 1% penicillin–streptomycin (Gibco), 1% GlutaMAX (Gibco), and 1% MEM non-essential amino acids (NEAA; Gibco). Baby hamster kidney fibroblasts expressing T7 polymerase (BHK-T7) were used for reverse genetics of SA11 RV, as described in Diebold et al. (Diebold et al., 2021). These cells were maintained in Glasgow’s Minimum Essential Medium (GMEM; Sigma) supplemented with 5% FBS, 10% tryptose phosphate broth (TPB; Sigma-Aldrich), 2% NEAA, 1% GlutaMAX, and 1% penicillin–streptomycin. All cell lines were maintained at 37°C in a humidified incubator with 5% CO₂.

### Plasmids

Plasmids encoding the complete genome of RV-SA11 (segments 1–11) were kindly provided by Dr. Takeshi Kobayashi. To generate the pT7/SClow plasmid, a gBlock (Integrated DNA Technologies) encoding the *SClow* open reading frame (ORF), flanked by a 5′ *AfeI* site and a 3′ *SacI* site, was synthesised and cloned into the pT7/NSP5 vector, replacing the wild-type *NSP5* ORF. The sequence of the pT7/SClow plasmid was confirmed by Sanger sequencing. Plasmids for transfection were prepared using a DNA Maxi Prep Kit (Qiagen) according to the manufacturer’s instructions, followed by ammonium acetate precipitation to obtain a final concentration exceeding 1 µg/µl.

### Reverse Genetics and Virus Propagation

BHK-T7 cells were seeded in 12-well plates to reach ∼70% confluency the following day. 24 hours later, a plasmid mixture was prepared in 250 µl pre-warmed Opti-MEM (Gibco), containing 0.8 µg each of pT7/VP1, VP2, VP3, VP4, VP6, VP7, NSP1, NSP3, and NSP4, 2.4 µg each of pT7/NSP2 and pT7/SClow, and 0.8 µg each of pcDNA3-NSP2 and pcDNA3-NSP5. To this, 34 µl TransIT-LT1 transfection reagent (Mirus Bio; Geneflow) was added. The mixture was incubated for 18 minutes at 25°C. BHK-T7 cell monolayers were washed with Eagle’s Minimum Essential Medium (Joklik modification, Sigma-Aldrich) supplemented with 1% GlutaMAX. The transfection mixture was then added dropwise, and cells were incubated at 37°C for 48 hours.

After 48 hours, confluent MA104 cells were harvested using trypsin-EDTA (Sigma-Aldrich), resuspended in DMEM containing 1 µg/ml trypsin, 1% GlutaMAX, and 1% NEAA, and overlaid onto the transfected BHK-T7 cells. Co-cultures were incubated at 37°C for 4 days. On day 7, cells were subjected to three freeze–thaw cycles to release virus particles. Cell lysates were clarified by centrifugation at 10,000 × g for 5 minutes, and the supernatants were treated with 2 µg/ml porcine trypsin at 37°C for 30 minutes. Trypsin-activated virus preparations were used to infect fresh monolayers of MA104 cells in FBS-free DMEM. Cells were incubated at 37°C until ∼80% cytopathic effect (CPE) was observed, at which point virus was harvested.

### Virus Propagation

For propagation, confluent MA104 cells were infected at an MOI of 0.01 using FBS-free DMEM supplemented with 0.5 µg/ml trypsin. Following infection, cells were lysed by three freeze–thaw cycles, and lysates were clarified by centrifugation at 10,000 × g for 5 minutes. For each passage, virus-containing supernatants were activated by incubation with 1 µg/ml trypsin at 37°C for 30 minutes prior to use.

To verify recombinant virus sequences, viral RNA was extracted from infected MA104 cells using the RNeasy Mini Kit (Qiagen), followed by reverse transcription with the SuperScript III Reverse Transcriptase Kit (Invitrogen). Amplified cDNA was generated using OneTaq Quick-Load 2X Master Mix (New England Biolabs) and analysed by Sanger sequencing.

### Determination of Viral Titres

Plaque assays were performed to quantify viral titres. MA104 cells seeded in 6-well plates were washed with FBS-free complete DMEM 30 minutes prior to infection. Serial tenfold dilutions of virus stocks (10⁻¹ to 10⁻⁸) were prepared in FBS-free medium and added to cell monolayers. Following a 2-hour incubation at 37°C, the inoculum was removed, and cells were overlaid with 1.2% Avicel (Sigma-Aldrich) in FBS-free DMEM containing 0.5 µg/ml trypsin. Plates were incubated at 37°C for 4 days. To fix the cells, the overlay was removed and replaced with 10% formaldehyde. Plates were incubated at 37°C for 1 hour, then stained with Coomassie Brilliant Blue for 1 hour at 37°C. Plaques were counted, and viral titres were calculated using the equation:

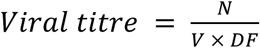

where *N* is the number of plaques, *V* is the volume of inoculum (ml), and *DF* is the dilution factor.

For TCID₅₀ assays, MA104 cells were seeded in 48-well plates (Sigma). Virus samples were serially diluted tenfold (10⁻¹ to 10⁻⁸) in serum-free DMEM. Each dilution was added to 8 replicate wells. After a 96-hour incubation at 37°C, wells were examined for cytopathic effect (CPE). TCID₅₀ values were calculated using the Reed–Muench method (REED and MUENCH, 1938) and expressed in plaque-forming units (PFU) per millilitre using the standard conversion (Pourianfar et al., 2012): PFU/ml = 0.7 × TCID₅₀/ml.

### Characterisation of Recombinant viruses Replication kinetics

Confluent MA104 cells (African green monkey kidney; ATCC CRL-2378.1) were infected in duplicate with either SA11-RV or SClow-RV at a multiplicity of infection (MOI) of 1. After a 60-min adsorption at 37 °C, the inoculum was removed, cells were washed once with 2 mM EGTA in PBS, and fresh serum-free DMEM containing 0.5 µg ml⁻¹ trypsin (Worthington) was added. Supernatants and cell lysates were harvested at 0, 4, 6, 8, 12, 16, 24 and 48 h post-infection (h p.i.). Virus was released by three freeze-thaw cycles (–80°C/37°C), clarified by centrifugation (16,000 ×g, 5 min), and the resulting supernatant was activated with trypsin (1 µg ml⁻¹, 30 min, 37 °C). Infectious titres were quantified in three independent plaque assays and expressed as mean plaque-forming units per millilitre ± SD.

### Genetic stability

To assess serial-passage stability, MA104 cells were infected with SClow-RV at MOI = 0.1 and passaged ten times (P1–P10). At complete cytopathic effect, cell culture medium was collected, clarified (16,000 ×g, 5 min), trypsin-activated as above, and titrated. Virus harvested from each passage served as inoculum for the next passage.

### Western blotting

RIPA buffer (150 mM NaCl, 1% Triton X-100, 0.1% sodium dodecyl sulfate, Tris-HCl pH-8, phosSTOP and cOmplete protease inhibitor cocktail (Roche)) was used to lyse RV-infected cells (MOI=20) at 4, 6, and 8 h.p.i.. Cell lysates were harvested and centrifuged at 15,000 rcf at 4 °C for 10 minutes. Supernatants were mixed with an equal amount of 2X Laemmli buffer (Bio-Rad) and heat-denatured at 98 °C for 10 min. Proteins were resolved by electrophoresis on 15% SDS polyacrylamide gels and transferred onto a nitrocellulose membrane (Millipore, Bedford, MA) using the Bio-Rad Trans-Blot Turbo Transfer System. Blots were then blocked with 5% (w/v) milk (MARVEL) dissolved in Phosphate buffered saline (PBS) buffer pH 7.4 (Oxoid). The membrane was incubated with primary antibodies (guinea pig anti-NSP5 (Papa et al., 2020) diluted 1:2500 in 5% (w/v) milk in PBS. The blots were next washed with 0.5% Tween-20 (Sigma-Aldrich) dissolved in PBS before incubation with anti-guinea pig IgG (H + L) cross-adsorbed secondary antibody DyLight 800 (1:10000; Invitrogen) and anti-actin hFAB rhodamine antibody (1:2500; Bio-Rad) in 5% milk (w/v) in PBS. The Bio-Rad ChemiDoc MP imaging system was used for fluorescent signal detection (DyLight800/Rhodamine filters). The band intensities of viral NSP5 were measured in Image Lab (version 6.1.0) and normalised to beta-actin band intensities and NSP5-specific antibody sensitivity toward WT-NSP5 and SC_low_ (400 ng WT-NSP5 band volume * 0.6687 = 400 ng SC_low_ band volume; simple linear regression). Statistical analysis was conducted using GraphPad Prism v10.

### Sample processing for transmission electron microscopy (TEM) imaging

MA104 cells were cultured in 35 mm ø Ibidi dishes with plastic coverslips (Ibidi-treat) and specimen processing for EM was carried out in these culture dishes. Confluent cells were infected in FBS-free medium with WT- or SC_low_-RVs at MOI=10. Samples were fixed at 8 h.p.i in fixative (2 % glutaraldehyde (TAAB) and 2 % formaldehyde (Merck) in 0.05 M sodium cacodylate buffer pH 7.4 (Sigma) containing 2 mM calcium chloride (Sigma)) overnight at 4°C. After washing 5x with 0.05 M sodium cacodylate buffer pH 7.4, samples were osmicated (1% osmium tetroxide (TAAB), 1.5 % potassium ferricyanide (AnalaR), 0.05 M sodium cacodylate buffer pH 7.4) for 3 days at 4°C. After washing 5x in DIW (deionised water), samples were treated with 0.1 % (w/v) thiocarbohydrazide (Merck) in DIW for 20 minutes at room temperature in the dark. After washing 5x in DIW, samples were osmicated a second time for 1 hour at RT (2% osmium tetroxide/DIW). After washing 5x in DIW, samples were blockstained with uranyl acetate (2 % uranyl acetate (Merck) in 0.05 M maleate buffer pH 5.5 (Merck)) for 3 days at 4°C. Samples were washed 5x in DIW and then dehydrated in a graded series of ethanol (50%/70%/95%/100%/100% dry) (Merck) and 100% dry acetonitrile (Macron), 3x in each for at least 5 min. Samples were infiltrated with a 50/50 mixture of 100% dry acetonitrile/Quetol (TAAB) resin (without BDMA) overnight, followed by 3 days in 100% Quetol (without BDMA). Then, the sample was infiltrated for 5 days in 100% Quetol resin with BDMA (TAAB), exchanging the resin each day. The Quetol resin mixture is: 12 g Quetol 651, 15.7 g NSA (TAAB), 5.7 g MNA (TAAB) and 0.5 g BDMA (TAAB). The Ibidi dishes were filled with resin to the rim, covered with a sheet of Aclar and cured at 60°C for 3 days. After curing, the Aclar sheets were removed and small sample blocks were cut from the Ibidi dish using a hacksaw. Thin sections (∼ 70 nm) were prepared using an ultramicrotome (Leica Ultracut E). Resin blocks were orientated with the cell-side towards the knife and sections were collected on bare 300 mesh copper grids (EM Resolutions) immediately when reaching the cell monolayer. Samples were imaged in a Tecnai G2 TEM (FEI/Thermo Fisher Scientific) run at 200 keV using a 20 µm objective aperture. Images were acquired using an ORCA HR high resolution CCD camera (Advanced Microscopy Techniques Corp, Danvers USA).

### Fluorescent Imaging Immunofluorescence

Cells were seeded in an µ-slide 8-well microscope slide (high glass bottom, Ibidi). At 90-100% confluency, cells were infected with RV. For immunofluorescence, cells were fixed with 4% Paraformaldehyde (PFA), permeabilised with 100mM glycine (Sigma-Aldrich) and 0.2% Triton X-100 consecutively. NSP5 was labelled with primary antibody GP anti-NSP5 (1:500), followed by secondary antibody goat anti-GP IgG (H + L) DyLight 550 (1:1000; Invitrogen). Cells were then stained with 300 nM 4’,6-diamidino-2-phenylindole (DAPI) for nucleus visualisation and imaged using the ONI Nanoimager S microscope connected to an Olympus 100x super apochromatic oil immersion objective (NA1.4). DAPI was imaged using a 405 nm laser at 19% power, while the NSP5-eGFP was detected with a 488 nm laser at 1% intensity. The fluorescence images of both live cells and fixed cells were acquired using a sCMOS camera (Andor Technology) with a pixel size of 0.117 µm. For quantification, automatic particle counting of viroplasms with sizes ranging from 0-10 µm^2^ (Yen) and nucleus with sizes ranging from 50-600 µm^2^ (Default) was performed in FIJI (Schindelin et al., 2012). Statistical analysis was performed using the GraphPad software (Prism v10).

### Single-molecule fluorescence in situ hybridisation (smFISH)

MA105 WT cells or MA104 NSP5 were seeded in an µ-slide 8-well microscope slide (high glass bottom, Ibidi). At 90-100% confluency, cells were infected with SC_low_-RV. For both cell lines, a mock-infected control was included. Cells were fixed at 8 h.p.i with 4% (v/v) methanol-free paraformaldehyde in nuclease-free PBS for 10 minutes. smFISH was conducted as described in (Strauss et al., 2023). RNA FISH probes are listed in https://cdn.elifesciences.org/articles/68670/elife-68670-supp1-v3.xlsx. Subsequent imaging of cells infected with SC_low_ RV was performed as described above. Imaging of cells infected with SC_low_ S67A RV was performed on a custom-built widefield microscope. The microscope frame (IX83, Olympus) is equipped with an LED light source (DC4100, Thorlabs) and a scientific complementary metal-oxide semiconductor camera (Zyla 4.2, Andor) controlled with the software Micro-Manager (Open Imaging). All images were acquired with a 60×/1.42 oil objective lens (PlanApoU, Olympus). Image analysis was performed as described above.

*In vitro* condensates were imaged using the ONI Nanoimager. Before imaging, samples were diluted in Phosphate buffered saline pH 7.4 (Merck KGaA). NSP2 was mixed with 0.5 µM NTA-Atto-488 (Merck KGaA) before addition of an NSP5 construct (either NSP5 SA11 nStrep, NSP5-HP SA11 nStrep, SC_low_ nStrep or SC_low_ HP nStrep, NSP5 IDR, NSP5 IDR HP or the NSP5 CTR) at an equimolar concentration to NSP2, to a working concentration of 25 µM per protein. 6 µL of this mixture were mounted on a Corning cover glass (CLS2980245; Merck KGaA, Darmstadt, Germany), which was placed on the objective using Olympus Low Auto Fluorescence Immersion Oil. Images were acquired by scanning N = 3 of a 50 - 500 µm x 80 - 800 µm region. Representative images were prepared using Fiji (Schindelin et al., 2012) Yen’s thresholding technique (Jui-Cheng Yen et al., 1995) allowed for subsequent automated particle analysis. Graphpad prism version 9.3.1 was used for creation of graphs and statistical analysis.

### Expression and purification of recombinant proteins

Recombinant NSP2 cHis (strain SA11 and strain RF) was expressed and purified as described previously (Schuck et al., 2001). Protein sequences were ordered as gBlocks from Integrated DNA technologies, carrying restriction sites for XbaI at the 5-prime end and for XhoI at the 3-prime end. Using restriction enzymes and T4 DNA ligase obtained from New England Biolabs, these gene fragments were cloned into a pET22 vector for subsequent expression in BL21(DE3) cells (New England Biolabs). NSP5 HP and SC_low_ HP carry S to D mutations at the following positions: 30, 37, 42, 56, 67, 101, 127 and 164. Expression and inclusion body isolation were conducted as described in (Borodavka et al., 2017), Inclusion body wash, solubilisation and dialysis as in (Martin et al., 2011). After dialysis, cleared supernatant was loaded onto a pre-equilibrated Strep-tactin superflow column (iba lifescience), and washed with 20 mM MOPS pH 7.1, 1 M NaCl for 20 column volumes (CV). Protein was eluted using 20 mM MOPS pH 7.1, 1 M NaCl, 2.5 mM Desthiobiotin for 5 CV. Protein-containing fractions were concentrated, and buffer was exchanged to 20 mM MOPS pH 7.1, 150 mM NaCl using an Amicon Spin column (Millipore) with a molecular weight cut-off of 3K. Protein concentrations were estimated spectrophotometrically using an extinction coefficient for NSP5 (HP) of 10,555 and SC_low_ (HP) of 19,160. The CTR peptide (YILDDSDSDDGKCKNCKYKRKYFALRMRMKQVAMQLIEDL) was obtained from the GenScript Biotech.

### Fluorescent labelling of recombinant proteins

NSP5 IDR and IDR HP were prepared for fluorescent labelling by incubating both protein samples in 10 mM tris(2-carboxyethyl)phosphine (TCEP) for 30 minutes at room temperature. TCEP concentration was reduced to 500 μM in 50 mM Tris-HCl pH 8.5 and 150 mM NaCl by using an Amicon Spin column (Millipore) with a molecular weight cut-off of 3K, and the samples were concentrated to a final volume of 250 μl each. A DyLight 488 Maleimide dye (Thermo Fisher Scientific) concentration of 1.25 mM was adjusted per sample. The labelling reaction was incubated for an hour at room temperature protected from light. After labelling, free dye was removed using a Slide-A-Lyze MINI Dialysis Device with a molecular weight cut-off of 3.5K (Thermo Fisher Scientific). Dialysis was performed against PBS pH 7.4 (Oxoid). Protein concentration and purity was monitored using a Spectrophotometer (Extinction coefficient IDR (HP): 10,555 and a correction factor (CF) of 0.147 for DyLight 488) and SDS-PAGE, respectively. The molar protein concentration was determined as follows: (C) = 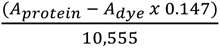

### Mass Photometry

Mass photometry measurements were conducted using a Refeyn Two^MP^ mass photometer and the Acquire^MP^ software (Refeyn Ltd, Oxford, UK, v 2023 R1.1) at 21°C at a 50 Hz frame rate and a 10.9 μm × 4.3 μm instrument field of view (46.3 μm^2^ detection area). Before 3 movies (n ≥ 3) of 60 s each was recorded per sample, protein stock was diluted in sterile-filtered phosphate buffered saline at pH 7.4 (806552-1L; Merck KGaA, Darmstadt, Germany). A ready-to-use sample carrier slide (Refeyn Ltd. Oxford, UK) was used in combination with a 6-well silicon sample cassette (Refeyn Ltd. Oxford, UK) for identification of the focal position (using 18 µl of buffer) and movie acquisition (by spiking the buffer with 2 µl of sample). The final sample volume was 20 µl with a working concentration of 100 nM. BSA, lgG and Thyroglobulin were used for contrast-to-mass calibration. The Discovery^MP^ software (Refeyn Ltd, Oxford, UK, v 2023 R1.2) was used for data processing, with a bin width of 5.8 kDa, and figure creation.

### PhaseScan Analysis

Microfluidic devices were formulated employing AutoCAD software and subsequently produced through traditional soft-photolithography techniques and Polydimethylsiloxane (PDMS)-on-glass devices (Arosio et al., 2016; Qin et al., 2010; Saar et al., 2017). Phase Scan was operated as described previously (Arter et al., 2022; Sneideris et al., 2023). Briefly, NSP2 was incubated with 5 µM Atto NTA-Atto-488 (39625; Merck KGaA, Darmstadt, Germany) and mixed with a version of NSP5 (HP), IDR (HP), SC_Low_ or SC_Low_ HP, supplemented with Alexa Fluor 647 dye (Thermo Fisher) at concentrations ranging from 2 µM to 25 µM in PBS pH 7.4. Protein mixtures at varying concentrations were encapsulated in water-in-oil droplets. An epifluorescence microscope (Cairn Research) equipped with a 10 × objective (Nikon CFI Plan Fluor 10×, NA 0.3) was used to take images of microfluidic droplets in the imaging chamber. An automated image analysis script was employed for the detection of condensates and the quantification of protein concentrations within individual droplets. The data was graphically represented as a scatter plot, accompanied by a colour-coded heat map overlay illustrating the estimated probability of phase separation (Sneideris et al., 2023). To determine the phase separation probability, the phase diagram was partitioned into grids with a bin size of 1 + 3.322 × log (total number of data points). Subsequently, the probability was calculated by dividing the total number of points labelled as phase-separated by the overall number of points within each grid (Sneideris et al., 2023).

### Hydrogen-Deuterium Exchange Mass-Spectrometry (HDX-MS)

HDX-MS experiments were performed using an automated HDX robot (LEAP technologies, USA) coupled to an Acquity M-class LC and HDX manager (Waters Corporation, UK). For NSP2 + NSP5 HP, a total of 5 µL of protein containing solution containing NSP2 (25 µM), NSP5 HP (25 µM), or a mixture of both (25 µM each) was added to 95 µL of deuterated buffer (1X PBS pD 7, in D_2_O). Samples were incubated at 4 °C for 0, 0.5, 5 and 10 min. The exchange reaction was quenched post labelling by addition of 75 µL of quench buffer (1X PBS pH 1.8, 0.1% DDM) to 75 µL of the sample. A total of 90 µL quenched sample was passed through an immobilised pepsin column (Enzymate, Waters Corporation, UK) and the resultant peptides were trapped on a VanGuard Pre-Column (Acquity UPLC BEH C18 [17 µm, 2.1 mm x 5 mm], Waters Corporation, UK) for 3 min. Separation of peptide fragments was achieved with a C18 column (75 µm x 150 mm, Waters Corporation, UK) eluting over a linear gradient of 0-40% (v/v) acetonitrile (0.1%[v/v] formic acid) in H_2_O (0.3% [v/v] formic acid) over 7 min at 40 µL min^-1^. Peptide fragments were detected using a Synapt G2-Si mass spectrometer (Waters Corporation, UK), operated in mobility-assisted data independent analysis with dynamic range extension enabled (HDMSe). Data was analysed using PLGS and DynamX software (Waters Corporation, UK). Pepsin was excluded from the analysis and restrictions for peptides in DynamX were minimum intensity = 1000, maximum sequence length = 25, minimum products per amino acid = 0.3, max ppm error = 10, file threshold = 2/3 replicates. Deuteros (Lau et al., 2019) was used to identify statistically significant protected and deprotected peptides with an applied confidence interval of 98%, visualised in Woods plots.

### Conservation Analysis

Sequences encoding NSP2 were selected as described in consensus sequence creation (Methods), and then aligned using the Clustal 2.1 tool (Madeira et al., 2022). Protein sequence conservation and subsequent model creation were performed using UCSF ChimeraX version 1.7 using the AL2CO entropy measure (Pei and Grishin, 2001). The conservation scores for NSP2 sequences were calculated using the and associated with the NSP2 SA11 structure 1L9V. NCBI IDs of NSP2 sequences used for analysis were: Q86505.1 (strain RF), AFK09596.1 (strain SA11), AKA40121.1 (S7_low_), AGF91999.1 (S6_low_), QIJ58153.1 (S5_low_), BBB18135.1 (S4_low_), ALJ83250.1, AHZ33501.1, QWY12027.1, AGE47301.1, ALJ83232.1, BBJ34822.1, ALQ56903.1, QWY12017.1, ACV73806.1, QCL12358.1, QCD16951.1, AAY55957.1, QWY12010.1, AGE46905.1, AGV31551.1, BAV57881.1, AGT98571.1, ADW11141.1, QCL12357.1 and AGE47337.1. The EMBL-EBI Job Dispatcher sequence analysis tools framework was used for multiple sequence alignment (Madeira et al., 2024), which was visualised using UCSF ChimeraX (Meng et al., 2023).

### Accession number(s)

Nucleotide sequences were deposited in GenBank and have the following accession numbers: pT7/ SC_low_, PP828582. The sequence for the whole plasmid pT7/ SC_low_ is available in Addgene (ID:220289).

## Acknowledgements

The authors thank C. Charalambos (University of Cambridge, UK) for plasmid and reagent preparation. We further thank the Cambridge Advanced Imaging Centre for sample processing for electron microscopy, and assistance during imaging. We thank Dr. Francesca van Tartwijk for help with microscopy imaging.

## Author contributions

JA: Conceptualisation, Data curation, formal analysis, investigation, methodology, validation, visualization, writing – original draft, writing – review & editing. XW: Data curation, formal analysis, investigation, methodology, validation, visualization, writing – original draft, writing – review & editing. DD: Data curation, formal analysis, investigation, methodology, software, validation, visualization, writing – original draft. TA : Formal analysis, investigation, methodology, validation, visualization. AC : Formal analysis, investigation, methodology, validation, visualization. CH: Formal analysis, investigation, validation, visualization. LS: Formal analysis, investigation, methodology, validation. KLS: methodology, software. RS: Formal analysis, investigation, methodology, validation, visualization. RM: Formal analysis, investigation. KS: Formal analysis, investigation. SHC: Formal analysis, investigation JDS: Conceptualisation, visualisation, writing – original draft, writing – review & editing. ANC: Conceptualisation, Supervision, funding acquisition. TPJK: Supervision, funding acquisition. AB: Conceptualisation, data curation, funding acquisition, methodology, project administration, resources, supervision, validation, writing – original draft, writing – review & editing.

